# A computational observer model of spatial contrast sensitivity: Effects of wavefront-based optics, cone mosaic structure, and inference engine

**DOI:** 10.1101/378323

**Authors:** Nicolas P. Cottaris, Haomiao Jiang, Xiaomao Ding, Brian A. Wandell, David H. Brainard

## Abstract

We present a computational observer model of the human spatial contrast sensitivity (CSF) function based on the Image Systems EngineeringTools for Biology (ISETBio) simulation framework. We demonstrate that ISETBio-derived CSFs agree well with CSFs derived using traditional ideal observer approaches, when the mosaic, optics, and inference engine are matched. Further simulations extend earlier work by considering more realistic cone mosaics, more recent measurements of human physiological optics, and the effect of varying the inference engine used to link visual representations to psy-chohysical performance. Relative to earlier calculations, our simulations show that the spatial structure of realistic cone mosaics reduces upper bounds on performance at low spatial frequencies, whereas realistic optics derived from modern wavefront measurements lead to increased upper bounds high spatial frequencies. Finally, we demonstrate that the type of inference engine used has a substantial effect on the absolute level of predicted performance. Indeed, the performance gap between an ideal observer with exact knowledge of the relevant signals and human observers is greatly reduced when the inference engine has to learn aspects of the visual task. ISETBio-derived estimates of stimulus representations at different stages along the visual pathway provide a powerful tool for computing the limits of human performance.

## Introduction

Newton’s work on the nature of light, some four centuries ago, initiated the quantitative understanding of vision. Since that time much has been learned about light, retinal image formation, fixational eye movements, and photon initiated excitations in the cone photoreceptors (Bowmaker, Dartnall, & Mollon, 1980; Wyszecki & Stiles, 1982; Rodieck, 1998; Engbert & Kliegl, 2004; Martinez-Conde, Macknik, & Hubel, 2004; Artal, 2015). Work continues to clarify how photoreceptor excitations are transformed into photocurrent and then to retinal and cortical signals that mediate visual perception (Baylor, Nunn, & Schnapf, 1984; Meister & Berry, 1999; Wandell, 1995; Pugh & Lamb, 2000; Angueyra & Rieke, 2013; Li et al., 2014).

All visual stimuli pass through the optics and retina, giving these structures a prominent role in defining the limits of vision. For example, the three-dimensional nature of human color vision can be understood in terms of the three types of cone photopigments that absorb light (Brindley, 1960; Wandell, 1995). Also, critical aspects of human pattern sensitivity depend on physiological optics (Robson, 1966; Campbell & Robson, 1968; Williams, 1985; Banks, Geisler, & Bennett, 1987). Quantification of human color and pattern sensitivity are critical for the imaging industry, including the design of cameras, displays, and printers; understanding the biological basis of visual sensitivity gives us confidence in the generality of the results and enables the diagnosis and targeted treatment of blinding disease.

Equally important, many aspects of visual perception are not explained by the initial stages of visual encoding. For example, human judgments of material appearance, the ability to recognize objects, and stereo vision depend on brain circuits that integrate information across space, time, and the two eyes. Attempts to understand these circuits can nonetheless benefit from a quantitative understanding of the initial encoding, as this determines the information available for perceptual inferences made by the brain.

Although our understanding of many properties of visual encoding may in principle be quantified using explicit computational models, putting such models to use in the practice of vision science is currently daunting. The relevant information is spread across a large literature, and integrating this information for a particular project typically requires a large effort. We developed the Image Systems Engineering Toolbox for Biology (ISETBio; http://isetbio.org) to make the relevant computations and data more accessible. ISETBio is an open-source software system that provides an image-computable model of the first stages of visual encoding.

Image-computable means that the calculations begin with a quantitative description of the visual image. An important special case supported by ISETBio is planar images presented on a computer display or the printed page. ISETBio also includes support for a more general case, in which the input is defined using a three-dimensional description that includes the location and shape of scene elements, as well as the spectral properties of each scene element. In this more general case, the retinal image is derived from the scene representation using ray-tracing methods (Pharr & Humphreys, 2010).

Computable methods are important because they can characterize visual representations for conditions that are beyond the reach of analytic formulations. When coupled with an inference engine that links the computed representations to performance on specific visual tasks, such as an ideal observer (De Vries, 1943; Rose, 1948; Tanner & Swets, 1954; Geisler, 1984, 1989), computable methods can (a) assess limits on performance, and (b) characterize the information available to brain circuits. We use the term *computational observer analysis* to describe image-computable methods combined with an inference engine (Lopez, Murray, & Goodenough, 1992).

This paper describes extensive updates to the previous versions of ISETBio (Farrell, Jiang, Winawer, Brainard, & Wandell, 2014; Jiang et al., 2017). We review and validate the updates by showing that the predictions for a computational observer implemented in ISETBio agree with analytical calculations of ideal observer pattern sensitivity derived in prior work (Banks et al., 1987; Geisler, 1984). We then explore how individual differences in human optics and cone mosaic impact predicted human performance. Finally, we analyze how performance varies with parameters and architecture of the inference engine. In particular, we consider inferences engines designed for pattern detection, and compare support vector machine (SVM) methods that learn about the stimulus with ideal observer methods where the stimulus and noise are known exactly.

Pattern sensitivity analysis is but one of many potential applications of ISETBio. We hope that making the software open-source and freely available will help others to develop analyses in new application areas.

## ISETBio overview

ISETBio computations are organized into a series of extensible methods that model the critical stages of visual encoding, from the visual scene through the optics, cone mosaic and inference engine (Figure 1). ISETBio scene methods represent the visual scene and enable calculations based on this representation. Scene methods include quantitative computer graphics calculations. These interoperate with optical image methods to compute retinal spectral irradiance from descriptions of relatively complex three-dimensional scenes. They also include methods for representing visual stimulicon-sisting of a spatio-temporal pattern specified as the spectral radiance emitted at each location and time on a flat screen. In this case, which is the one we use in this paper, the scene specification can be in terms of RGB values and is coupled to display calibration data, most importantly the spectral radiance of each of the display primaries (Figure 1A). The ability to represent stimuli presented as images on a flat display is important for modeling many psychophysical experiments. We have also implemented methods that support modeling of psychophysical stimuli presented in Maxwellian view (Tuten et al., 2018).

**Figure 1:**
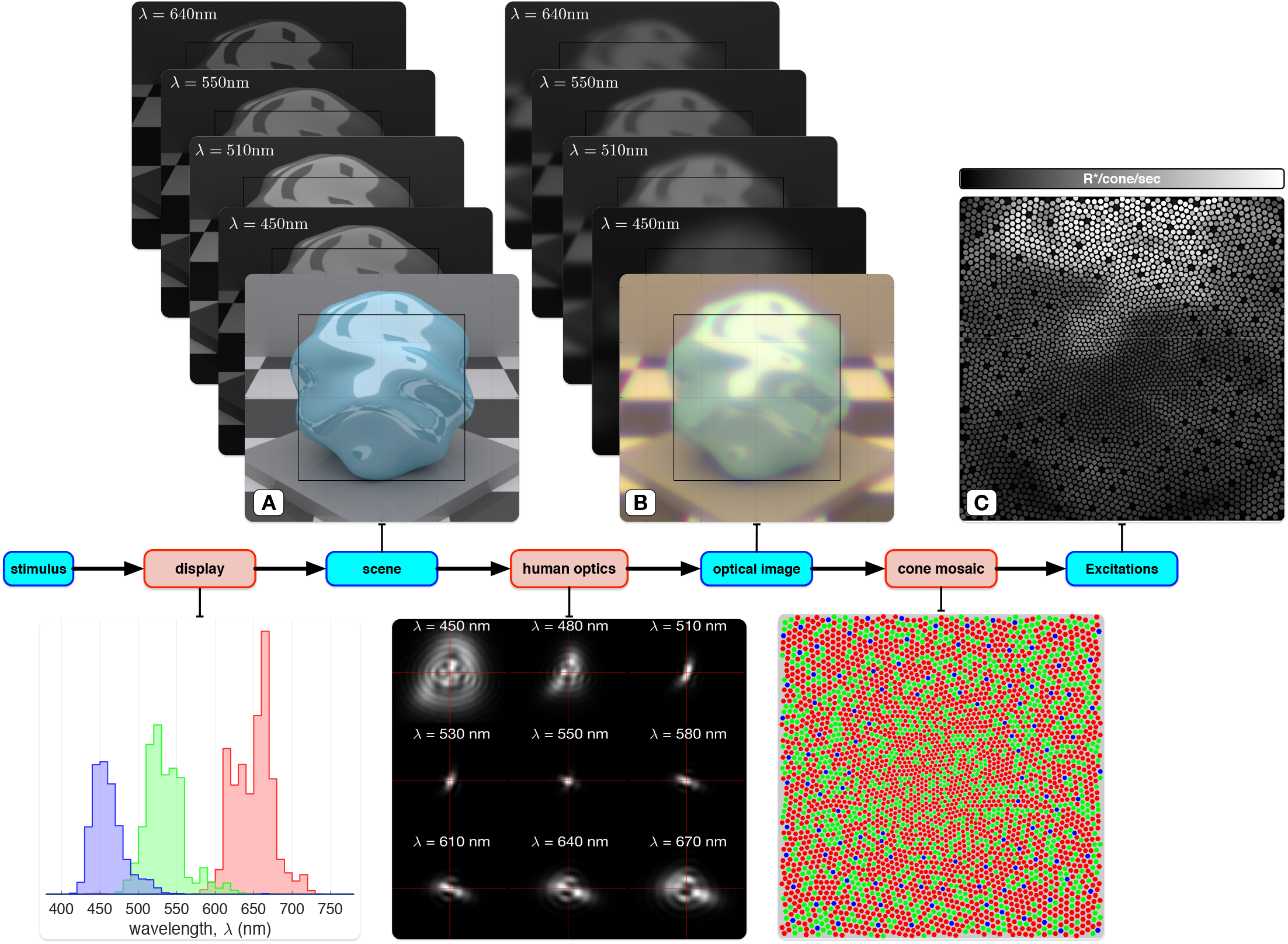
Flowchart of computation in ISETBio. **A**. The visual stimulus is represented as an ISETBio scene, in this case an image on a display. An ISETbio scene represents the scene radiance at a number of wavelengths. Here, the spectral power distributions of the display primaries (lower portion of the figure) and the pixel spatial sampling are used to convert stimulus RGB values to the spatial-spectral radiance. An RGB rendition of the scene is depicted in front of the spectral radiance stack. In the calculations reported in this paper, wavelengths are sampled between 380 nm and 780 nm at 5 nm spacing. In the image a subset of the sample wavelengths is shown. **B.** ISETBio optical image methods transform the scene to the retinal spectral irradiance. These methods blur the scene radiance image with a set of wavelength-dependent point spread functions (example for one individual shown in the lower portion of the figure). The methods also account for spectral transmission through the lens. Spectral transmission through the macular pigment is handled as part of the computation of cone excitations. **C**. ISETBio cone mosaic methods compute cone excitation rates, which are indicated by grayscale in the figure. The S cones appear black; they are excited less than the L and M cones because of selective absorption of short wavelength light by the ocular media. In the mosaic shown (lower image), cone density decreases and cone aperture increases with eccentricity, and there is a central S cone free region.

The spectral irradiance incident at the retina is calculated from the scene representation using ISETBio optical image methods (Figure 1B). This computation accounts for critical physiological optics factors, including pupil size, wavelength-dependent blur, and wavelength dependent transmission through the crystalline lens. The optical image calculations used in this paper account for the wavefront aberrations of the eye’s optics, measured using a wavefront-sensor (Thibos, Hong, Bradley, & Cheng, 2002) and specified by Zernike polynomial coefficients. These measurements determine a set of wavelength-dependent point spread functions.

The spatial pattern of cone excitations is computed from the retinal irradiance using ISETBio cone mosaic methods. These methods transform the spectral irradiance at the retina into cone excitations (Figure 1C). The cone mosaic methods include parameters which control factors such as (a) the relative number of L, M and S cones, (b) the existence and size of an S-cone free zone in the central fovea, (c) the cone spacing, inner segment aperture size and outer segment length as a function of eccentricity, (d) the macular pigment density, and (e) the cone photopigment density. These parameters all affect the number of cone excitations.

In a contrast sensitivity experiment, the subject discriminates between a spatially-uniform pattern (null stimulus) and a cosinusoidal grating pattern (test stimulus). In ISETBio, we use the term inference engine to describe methods that link the computed visual system responses to psychophysical performance. Inference engine methods make decisions in a simulated visual task, based on the stimulus representation at different processing stages along the visual pathway. Figures 2A and 2B depict the retinal image contrasts seen by the L, M and S cones, for the null stimulus and for a 16 c/deg grating test stimulus with 100% Michelson contrast. Note that, for this stimulus, aberrations reduce L-and M-cone retinal image contrast by a factor of 2 relative to the stimulus image, whereas the retinal S-cone contrast is reduced by a factor of 10. Aberrations also shift the spatial phase of the S-cone contrast with respect to that of L- and M-cone contrasts (Figure 2B).

**Figure 2:**
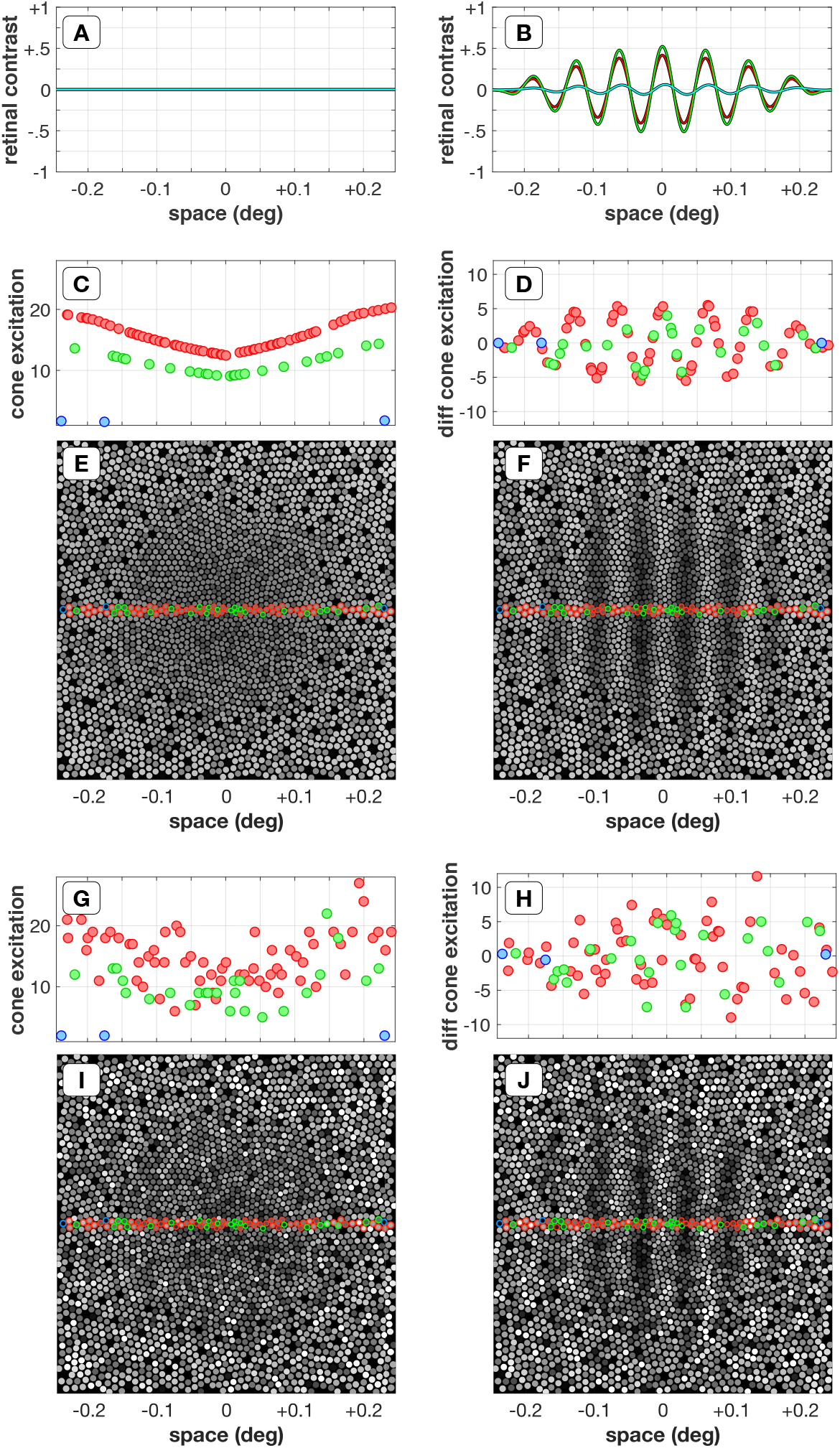
Stimulus representations in ISETbio. Representations of a uniform field (null stimulus) and of a 16 c/deg, 100% Michelson contrast cosinusoidal grating (test stimulus) are depicted in the left and right columns, respectively. **A&B**. Retinal contrast along the horizontal meridian for a null and test stimulus, respectively. These spatial contrasts are depicted as seen by the L-cones (red), M-cones (green) and S-cones (cyan). **C&D**. Mean cone excitation (number of photon absorption events within a 5 msec time bin) for cones along the horizontal meridian to the null stimulus and mean differential (test-null) cone excitation to the test stimulus, respectively. **E&F**. Mean cone excitation pattern of the entire mosaic to the null and the test stimulus. **G&H**. A single excitation instance of cones along the horizontal meridian to the null stimulus and a single differential excitation instance to the test stimulus, respectively. **I&J**. A single excitation instance of the entire mosaic to the null and the test stimulus.

Figures 2C and 2D plot the excitation level of cones along the horizontal meridian for the null stimulus, and the differential excitation for the test stimulus. The mean cone excitation increases with eccentricity because of changes in cone aperture with eccentricity. The mean excitation of the entire cone mosaic is depicted in Figures 2E and 2F. Figures 2G and 2H also depict the stimulus representation at the cone mosaic, but for a single response instance in which Poisson noise is added to the mean excitations. It would be challenging to discriminate between the two stimuli by looking at single response instances of just a few cones. Spatial integration across the cone mosaic will improve performance, as can be appreciated by visual comparison of Figures 2I and 2J. The figures illustrate the cone excitations to a supra-threshold 100% contrast grating, not to a grating at contrast threshold, and ultimately classification performance cannot exceed the limits imposed by the Poisson noise inherent in these excitations.

In this paper, we consider inference engines that model a two-alternative forced choice version of the contrast sensitivity experiment, and we use response instances at the level of cone mosaic excitations to predict the probability of correct discrimination between gratings and a uniform field. Performance is limited according to how well the inference engine is matched to the task (the classifier’s calculation efficiency; Barlow, 1964; Pelli, 1990), as well as the difference between the representations of the stimuli relative to those of noise, i.e., trial-by-trial fluctuations in the representations. As noted above, Poisson noise is inherent to cone excitations and is a critical limiting factor for performance at this stage of encoding.

## Results

### Pattern sensitivity validation

Complex software requires explicit testing of (a) the individual components (unit testing), (b) component communication (integration testing), and (c) the overall system (validation). The ISETBio software includes a number of such tests, as well as methods to check that new software methods do not invalidate previously established tests (regression testing). In this section we describe validation testing of a complex computation that utilizes key ISETBio methods. We show that the ISETBio implementation - including stimulus definition, physiological optics, and cone excitations - matches a precise analytical calculation for an ideal observer’s contrast sensitivity to known spatial harmonic patterns (signal-known-exactly) by Banks et al. (1987). This test is designed to provide confidence in the basic implementation and the validity of the subsequent explorations of how physiological optics, the cone mosaic, and the inference engine influence human pattern sensitivity.

We computed ISETBIO contrast sensitivity functions (CSFs) using parameters that match those used by Banks et al. (1987). These included a 2 mm pupil diameter, a point spread function (PSF) derived from early line spread function (LSF) measurements (Campbell & Gubisch, 1966; Geisler, 1984), a regularly-spaced hexagonal cone mosaic comprising an approximation of L and M cones in a 2:1 ratio, cone center-to-center spacing of 3 *μ*m and a cone inner segment aperture of 3 *μ*m. There was one small difference within the cone mosaic. Banks et al. (1987) calculated for a mosaic in which all cones were of the same type, each with a luminance spectral responsivity (2L+M); we modeled a mosaic consisting of distinct L and M cones in a 2:1 ratio.

Performance (probability correct) was estimated for each grating contrast and spatial frequency separately. We simulated a two-alternative forced-choice task, using an ideal observer classifier that selects which of the two alternatives (test–null or null–test) was more likely to generate the observed cone excitations. The test stimulus was a spatial grating of known contrast, frequency, and position; the null stimulus was a spatially uniform field. The simulated duration of the test and null stimulus was 100 msec on each trial, with the stimuli presented in random order. The ideal observer’s percent correct was calculated analytically given knowledge of the mean number of excitations (within 5 msec time bins over the 100 msec stimulus duration) of each cone to each stimulus and the assumption of Poisson noise. We fit the psychometric function (percent correct as a function of stimulus contrast) for each spatial pattern with a cumulative Weibull (Kingdom & Prins, 2010, http://www.paiamedestooibox.org). We took threshold to be the stimulus contrast corresponding to 70.71% correct. Sensitivity is the reciprocal of threshold contrast.

Figure 3A compares CSFs for three mean luminance levels (3.4, 34, and 340 cd/m). The solid lines show the ideal observer CSFs obtained by Banks et al. (1987), digitized from their Figure 2. The corresponding points show the CSFs computed using ISETBio with its implementation of the ideal observer inference engine. The ISETBio computation agrees well with that of Banks et al. (1987) across all spatial frequencies and luminance levels. In addition, the sensitivity ratios between the different mean luminance levels (bottom panel of Figure 3A) cluster around the ratios (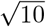 and 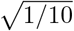) expected from the square-root law for Poisson-limited sensitivity (De Vries, 1943; Rose, 1948). We take this agreement as an important system validation of the ISETBio implementation.

**Figure 3:**
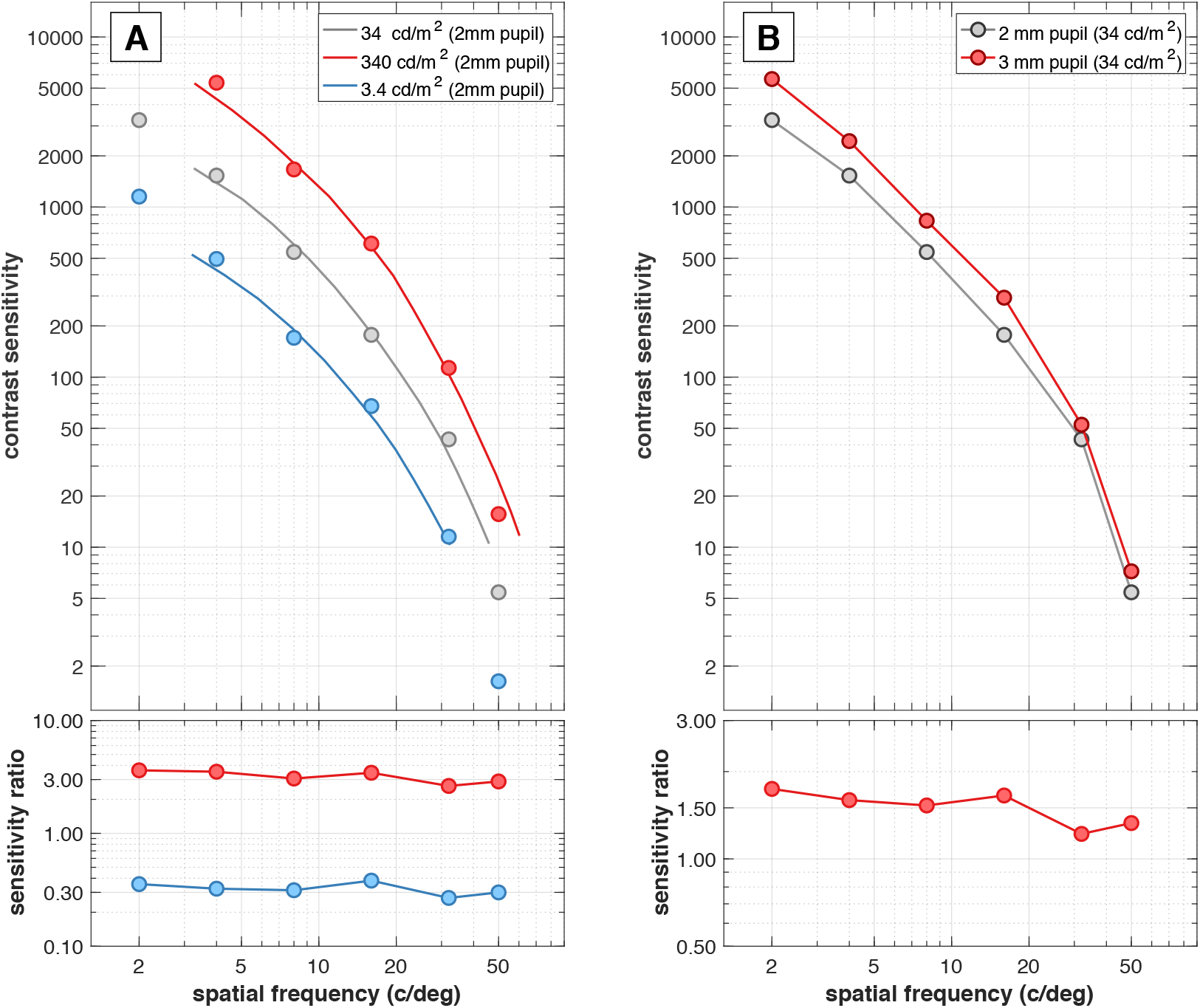
ISETBio validation. **A**. The top plot depicts the contrast sensitivity functions (CSFs) for a 2 mm pupil diameter and three mean luminance levels. Solid lines show ideal observer CSFs, digitized from Banks et al. (1987). Symbols show CSF values calculated using ISETBio with matched parameters. The bottom plot depicts the ratio of contrast sensitivities between the 3.4 and 34 cd/m mean luminances (blue) and between the 340 and 34 cd/m mean luminances (red). **B**. Top plot depicts the ISETBio ideal observer CSFs compared for 3 and 2 mm pupil diameters. Other parameters matched Banks et al. (1987). In particular the optical PSF was held constant across this comparison. The bottom plot shows the ratio of the 3 mm and 2 mm contrast sensitivities.

As a further check, we assessed the impact of increasing pupil diameter on the CSFs computed using ISETBio (Figure 3B). A 2 mm pupil is used for the comparison with calculations and psychophysical data reported by Banks et al. (1987), as their data were collected using a 2 mm artificial pupil. This size is smaller than we expect for natural viewing of the stimuli. When a 30 year old views a 50 deg, 34 cd/m adapting field binocularly the expected pupil diameter is 3.4 mm (Watson & Yellott, 2012); when the adapting luminance is 100 cd/m the expected pupil diameter is 3.0 mm. For Poisson signals, sensitivity should increase with the square root of retinal irradiance, and retinal irradiance is proportional to the square of pupil diameter. Thus we expect the sensitivity ratios for the 3-mm vs. the 2-mm CSF to be 1.5 across all spatial frequencies. This is confirmed to good approximation, which further validates the software implementation. Note that we did not change the optical point spread function for this test, although a change would be expected in a simulation aimed at fully understanding the impact of a change in pupil size.

Taken together, these computations ground the ISETBio ideal observer implementation in the analytical literature and validate the use of ISETBio for exploring how changes in visual system parameters impact the estimated upper bound for the spatial CSF.

### Cone mosaic

The Banks et al. (1987) psychophysical data were collected using a constant number of grating cycles across changes in spatial frequency. Thus the spatial extent of the stimuli was larger for lower spatial frequencies. Banks et al. (1987) employed a constant-density cone mosaic. For the human retina, however, cone density declines as a function of eccentricity; this decline is particularly rapid across the central fovea (Curcio, Sloan, Kalina, & Hendrickson, 1990). To explore how a change in cone density affects the spatial CSF for constant cycle stimuli, we developed new methods to implement realistic cone mosaics (outlined in section Cone mosaics). These cone mosaic methods retain the approximately hexagonal cone packing of central retina while decreasing the cone density with eccentricity.

We compared the ideal observer CSF calculations for the regularly-spaced hexagonal L/M cone mosaic with 3.0 *μ*m inner segment diameter used by Banks et al. (1987), with three different eccentricity-dependent mosaics. For these calculations, we simulated a mean stimulus luminance of 34 cd/m^2^ and a 3 mm pupil. The optical PSF matched the one used by Banks et al. (1987). The Banks et al. (1987) mosaic is illustrated in Figure 4A. It contains only L and M cones in a 2:1 ratio, with hexagonal packing at 3 *μ*m cone spacing and a 3 *μ*m cone inner segment diameter. The eccentricity-dependent cone mosaics are illustrated in Figure 4C-4D. The first also comprised only L and M cones (2:1 ratio, Figure 4B). In this mosaic, cone density decreases according to the measurements of (Curcio et al., 1990), and cone inner segment diameters were modeled at 1.6 *μ*m. Another eccentricity-dependent cone mosaic included L, M and S cones in the ratio 0.62:0.31:0.07, with S-cones starting to appear at eccentricities > 0.1 deg (Figure 4C). For the final eccentricity-dependent mosaic we further modeled eccentricity-dependent changes in the cone inner segment diameters and outer segment lengths (Figure 4D). Both of these factors affect cone excitation efficiency. Additional details are provided in the Eccentricity-dependent cone efficiency correction section. Figure 4E shows the effect of cone mosaic on the ideal observer CSF. The CSF plotted in gray replots the simulation of the Banks et al. (1987) constant-density mosaic from Figure 3B. The CSFs plotted in red, blue and green show the CSFs obtained with the three eccentricity-dependent mosaics. At low spatial frequencies these deviate systematically from the CSF of the constant-density mosaic. The size of the deviation is quantified in the sensitivity ratio plots in the bottom panel.

**Figure 4:**
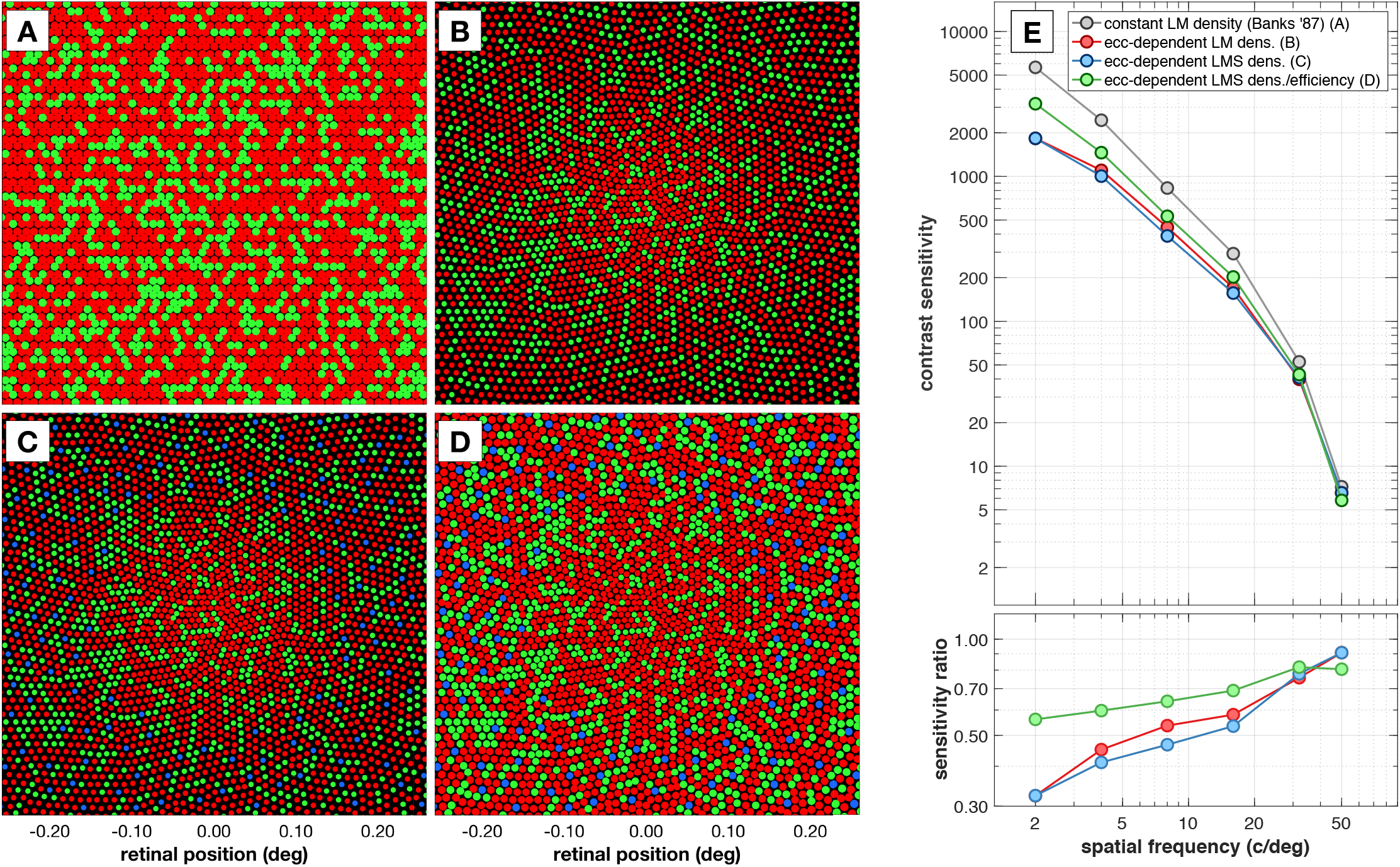
Effects of cone mosaic. **A-D**. Central 0.5 × 0.5 degrees of the employed mosaics. **A**. The mosaic used by Banks et al. (1987) has a regular hexagonal cone packing with 3 *μ*m cone spacing, 3 *μ*m cone aperture, and each cone has a luminance spectral sensitivity. We replicated the Banks et al. (1987) results using a regular mosaic with randomly assigned L and M cones in a 2:1 ratio. **B**. A mosaic with eccentricity-dependent cone density with only L and M cones in a 2:1 ratio. This mosaic has fixed cone inner segment diameter and outer segment length, e.g., independent of eccentricity. **C**. A mosaic with eccentricity-dependent cone density and fixed cone inner segment diameter and outer segment length with L, M and S cones. **D**. A mosaic with eccentricity-dependent cone density and cone inner segment diameter/outer segment length, also with L, M and S cones. In **B,C,D**, cones in the central region are separated by 2 *μ*m, which corresponds to the 250,000 cones/mm peak cone density reported by Curcio et al. (1990). The aperture to cone spacing ratio (0.79) is close to the 0.82 value suggested by Miller and Bernard (1983), and Curcio et al. (1990). In **C,D** the L:M:S cone ratios are 0.62:0.31:0.07 with an S-cone free central region. S-cone spacing outside of this central region was constrained to be relatively regular. E. Contrast sensitivity functions for the four mosaics in **A-D**, computed for a 3 mm pupil. Calculations were done using a 3 mm pupil diameter and the same PSF employed by Banks et al. (1987).

The effect of mosaic density is most pronounced for the two mosaics with constant inner segment diameter and outer segment length. For these mosaics, the drop in relative sensitivity occurs because in the constant cycles paradigm, low frequency stimuli extend further into the periphery where cone density is lower. This leads to lower total cone excitations in response to the stimuli compared to the constant-density mosaic, and thus lower ideal observer sensitivity. The addition of S cones has a negligible effect.

The drop in sensitivity for the low spatial frequencies is mitigated for the mosaic that includes a space-varying change in cone inner segment diameter and outer segment length. The net effect of the change in these factors is to increase the number of excitations per cone as eccentricity increases, partially offsetting the reduction in total excitations caused by reduction in cone density. Even for this mosaic, however, there is a notable decrease in low spatial frequency sensitivity compared to the constant density mosaic CSF. For the remaining calculations, we use the eccentricity-dependent mosaic shown in Figure 4D.

### Optics

There have been significant improvements in the ability to measure the optical quality of the eye since early measurements of the human line spread function (Westheimer & Campbell, 1962; Campbell & Gubisch, 1966). In particular, wavefront aberration measurements in individual human eyes (Liang & Williams, 1997) enable calculation of the corresponding point spread functions (PSFs); (Thibos, Ye, Zhang, & Bradley, 1992; Goodman, 2005; Watson, 2015). We examined how wavefront aberration based PSFs affect the derived CSF and contrasted this to the CSF derived by Banks et al. (1987) based on work from Geisler (1984). To do so we used the Thibos et al. (1992) data set, which includes a good sample of on-axis wavefront aberration measurements. However, as Thibos et al. (1992) point out direct averaging of the PSFs (or of the Zernike polynomial coefficients) results in a point spread function that differs qualitatively from any of the underlying measurements: the averaging process removes the idiosyncratic PSF structure found in most eyes. In addition, the optical modulation transfer function (MTF; absolute value of the complex OTF) obtained from the average of the individual eye Zernike coefficients is sharper than the average of the optical MTFs obtained from the same set of coefficients. This happens because the mean Zernike coefficient for defocus is near zero; some subjects have positive defocus while others have negative defocus, and the resulting MTF represents optics that are sharper than typical. Given these issues, we decided to derive CSFs based on PSFs from 5 subjects which were selected to cover the range of PSFs reported by Thibos et al. (2002). The selection process is described in detail in the Selecting representative Thibos subjects section.

Figure 5A depict the PSF used by Banks et al. (1987), and Figures 5B-5F depict the PSFs (at 550 nm) of the five subjects selected from the Thibos data set. All the PSFs were computed assuming a 3 mm pupil. Note that the PSF used by Banks et al. (1987) has no dependence on wavelength, whereas the wavefront-derived PSFs account for both higher order aberrations and longitudinal chromatic aberration. Figure 5G compares the ideal observer CSF obtained with the PSF used by Banks et al. (1987) with ideal observer CSFs obtained using optics of the five selected Thibos subjects. All CSFs agree well at low spatial frequencies, but the wavefront-derived functions fall off less rapidly than the Banks et al. (1987) CSF for spatial frequencies above 5 c/deg. This difference is substantial at frequencies above 30 c/deg, approaching a factor of 5 at 60 c/deg. The higher sensitivity arises because the wavefront-derived PSFs (Figures 5B-5F) are somewhat narrower than the PSF used by Banks et al. (1987) (Figure 5A). These results also show that variations in optics may lead to considerable individual variation in the CSF at high spatial frequencies. In summary, CSFs derived based on modern measurements suggest that typical observer optics enable a higher sensitivity at high spatial frequencies than the Banks et al. (1987) estimate. We selected the PSF of Subject 3 as a “typical” human PSF. All calculations from this point on were conducted using that PSF.

**Figure 5:**
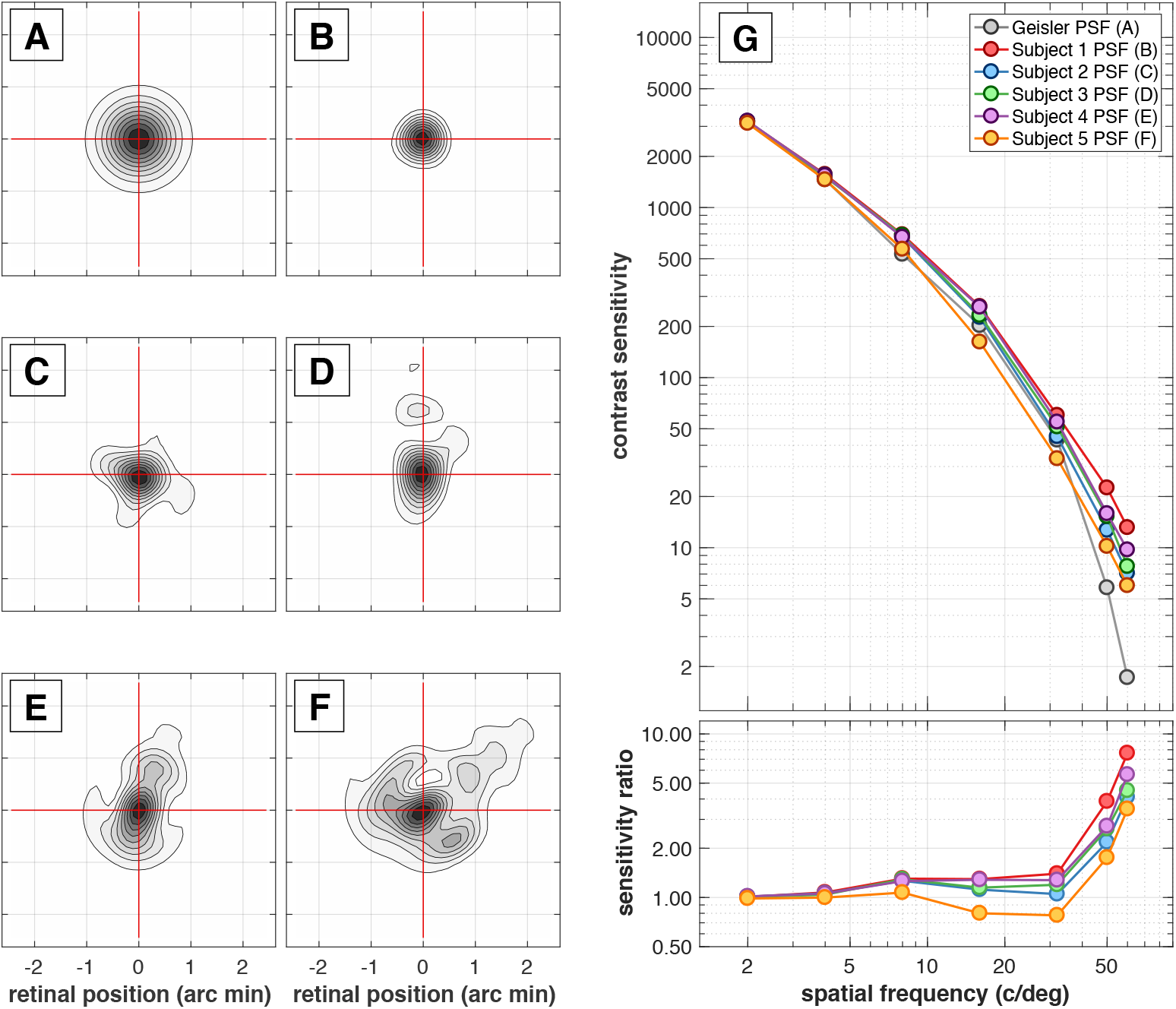
Effects of optics. **A-F**. Contour plots of the employed PSFs at 550 nm. Note that the Banks et al. (1987) PSF, displayed in **A**, is identical across all wavelengths, whereas the wavefront-based PSFs, displayed in **B-F**, vary with wavelength. We take the PSF of Subject 3 (depicted in **D**) to represent typical human optical quality. **G**. Contrast sensitivity functions for five individual point spread functions derived from wavefront aberration data reported by Thibos et al. (2002), compared with the CSF obtained using the PSF employed by Banks et al. (1987). For these calculations we simulated the eccentricity-dependent density and efficiency LMS cone mosaic.

### Inference engine

The ideal observer calculations reported thus far characterize the information available in the mosaic excitations when the spatiotemporal dynamics of the mean response and the statistics of the noise for the test and null stimuli are known exactly. This analysis provides an upper bound on performance, but the signal-known-exactly assumption is unlikely to match the computations of the neural mechanisms that process the cone mosaic signals.

An inference engine based on a Support Vector Machine (SVM) classifier (Scholkopf & Smola, 2002; Manning, Raghavean, & Schutze, 2008) uses training data to learn the parameters of a linear classifier. This classifier separates the visual representations of the test and null stimuli. Here the visual representation is at the cone excitations and trial-to-trial variability is due to Poisson noise, and we have focused on that case. Variability can also arise due to other factors, such as fixational eye movements, fluctuations in pupil size and accommodation, and noise in the neural representation at sites central to the cone excitations.

Figure 6 illustrates the idea underlying the SVM-based inference engines in the context of our two-alternative forced choice paradigm. Two scenes, one specifying the test stimulus, *S_t_*, and one specifying the null stimulus, *S_n_*, are processed through the ISETBio simulation pipeline. N response instances are computed for each of the test and null stimulus, 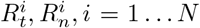. The samples for each stimulus differ because of Poisson noise. The sample data are divided into two sets, one used for training and another used for evaluation (held out data). Response vectors are formed comprising null and test excitations in the order of the two possible types of trials (test–null or null–test). For computational efficiency, a dimensionality reduction algorithm may be used to extract a low-dimensional representation of these responses; two-dimensions are illustrated in Figure 6 (red and blue data points). A linear SVM classifier is trained to derive a separating hyperplane (black line) which maximizes the separation between the two types of trials, and the classifier’s accuracy is evaluated on the held out data. The entire procedure is repeated for a range of stimulus contrasts, defining a psychometric function (percent correct as a function of stimulus contrast, gray points). A cumulative Weibull function is fit to the data, and the contrast level at 70.71% correct is considered the threshold.

**Figure 6:**
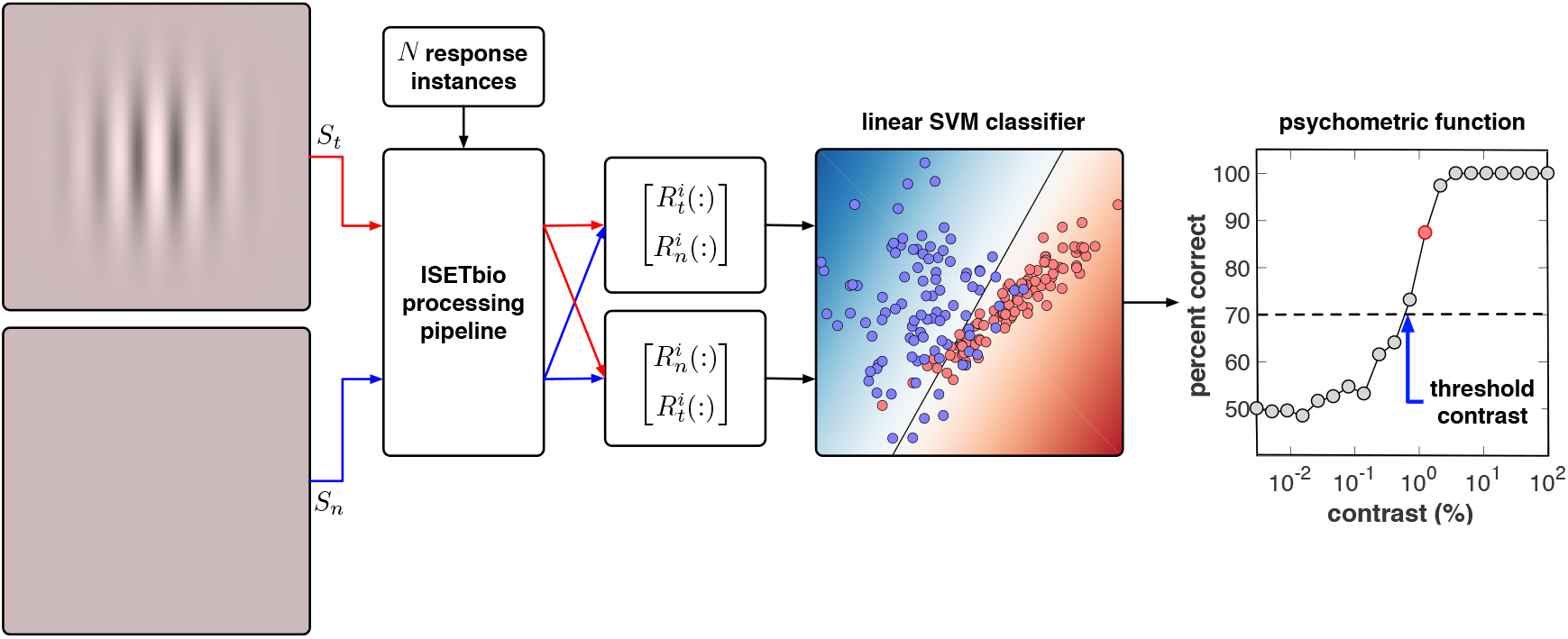
Illustration of SYM-based inference engine. Scenes describing the test stimulus, *S_t_*, (top left) and the null stimulus, *S_n_* (bottom left) are constructed. Each is run through the ISETBio pipeline multiple times to produce *N* instances of cone mosaic responses to each stimulus, 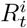 and 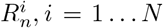. Each response instance includes an independent draw of Poisson isomerization noise. To simulate a two-alternative forced choice paradigm, composite response vectors are formed, with the response component to the test stimulus followed by the response component to the null stimulus, and vice versa. A dimensionality reduction algorithm may be used to extract a low-dimensional feature set from these composite responses; in this illustrative example, a two-dimensional set is shown. The data are divided into training and evaluation sets. The training set is used to train a linear classifier which learns the parameters of a hyperplane (shown as black line) that optimally separates instances of the two stimulus (null–test and test–null) orders (red and blue data points). The performance of the classifier is then obtained on the evaluation set. This process is repeated for a series of stimulus contrasts, leading to a simulated psychometric function, from which threshold is extracted. The red point in the plotted psychometric function shows performance for the classifier illustrated in the figure. Threshold is taken as the contrast that corresponds to 70.71% correct, based on a fit to the simulated psychometric function.

We analyzed the sensitivity of the ideal observer to that of two different SVM-based computational observer inference engines. The first SVM-based inference engine reduces the dimensionality of the signals in the full cone mosaic to 60 by projecting response vectors to the space of the 60 principal components derived from the entire date set (SVM-PCA engine). The second SVM-based inference engine reduces the dimensionality of the signals in the full cone mosaic to 20, the number of 5 msec time bins within the 100 msec presentation time, by taking the inner product of the mosaic response at each time bin with a spatial pooling template. The template used for each spatial frequency was derived from the contrast profile of the test stimulus at that spatial frequency as described in section Inference engine (Figure 13). These two SVM-based inference engines differ in how much must be learned from the training set. The SVM-PCA inference engine is provided with no a priori information about the stimuli, while the SVM-Template inference engine, like the ideal observer inference engine, has perfect knowledge of the stimuli. Unlike the ideal observer, however, the SVM-Template inference engine is not provided with knowledge about the structure of the noise, nor of the optimal criterion to apply to the underlying decision variable.

Figure 7A illustrates effects of the inference engine. In these simulations we used a 3 mm pupil, the typical human PSF (Subject 3 of Figure 5) and the cone mosaic with eccentricity-dependent density and efficiency (Figure 4D). We make three observations. First, as expected, the performance of both SVM classifiers is worse than the performance of the ideal observer, with CSF ratios that vary between 0.1 and 0.35 across the spatial frequency range (Figure 7A, bottom panels). Second, the ratio of performances of the SVM-Template classifier to the ideal observer inference engine is roughly constant with spatial frequency, whereas the ratio for the SVM-PCA inference engine varies strongly with spatial frequency. Third, the performance of the SVM-PCA inference engine is worse than that of the SVM-Template inference engine for low spatial frequencies, but better for spatial frequencies above 16 c/deg.

**Figure 7:**
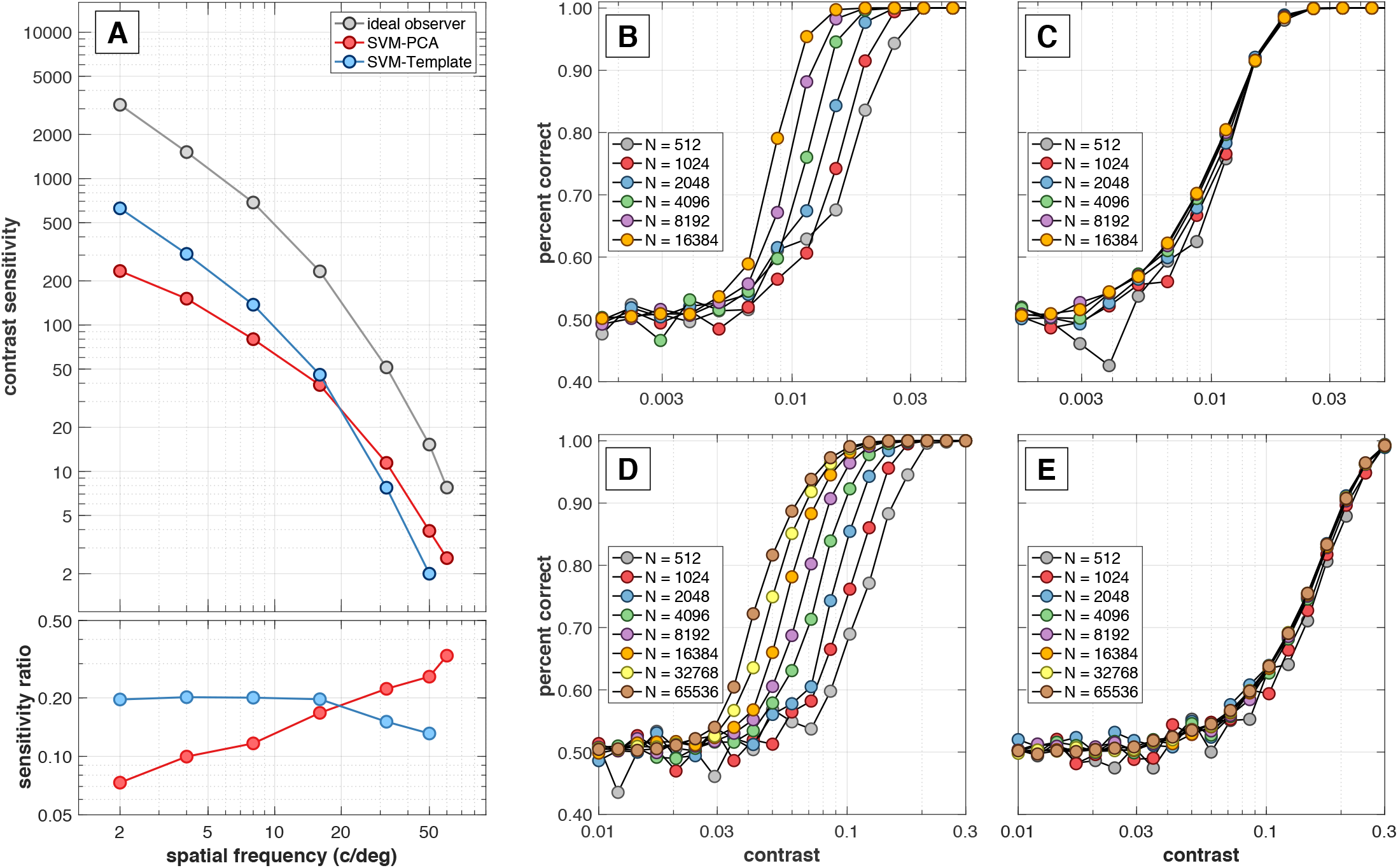
Effect of inference engine and training set size. **A**. Effects of different inference engines on the CSF. We used the typical subject PSF (Subject 3 from Figure 5 / Table 1), the eccentricity-dependent cone density and cone efficiency LMS mosaic and a data set consisting of 1024 response instances. Note that both SVM-based inference engines are about 5 times less sensitive than the ideal observer signal-known-exactly inference engine. **B&C**. Psychometric functions for the SVM-PCA and SVM-Template inference engines, respectively, for the 8 c/deg stimulus computed using training data sets of 512-16,384 instances. **D&E**. Psychometric functions for the SVM-PCA and SVM-Template inference engines, respectively, for the 32 c/deg stimulus, computed using training data sets of 512-65,536 instances.

The spatial frequency dependence of SVM-PCA inference engine performance may be due to an interaction between stimulus dimensionality and learning. At low spatial frequencies the activated mosaics are large and the response vectors have a high dimensionality. In this case the SVM-PCA classifier might be inefficient when we use only 1024 response instances to extract the principal components and to train the SVM. On the other hand, 1024 instances of the smaller dimensionality responses to higher spatial frequency stimuli appears sufficient to train a good classifier, resulting in a relative increase in performance with spatial frequency. We suspect that the performance of the SVM-Template inference engine is approximately constant with spatial frequency, relative to the ideal observer inference engine, because the SVM-Template is provided with information about the spatial structure of the stimuli.

To investigate further, we examined performance of the different inference engines as a function of the size of the training set for 2 spatial frequencies, 8 and 32 c/deg. The psychometric curves of the SVM-PCA inference engine depend strongly on the data set size, shifting to the left as the data set size increases (Figures 7B and 7D). The SVM-Template inference engine performance is relatively stable, changing only slightly with the size of the training set (Figures 7C and 7E).

Figure 8 quantifies the effect of training data set size on the computed contrast sensitivity for three stimuli (8, 16, and 32 c/deg), and for the ideal observer and the two SVM-based computational observers. Ideal observer performance does not depend on the number of trials because its performance is computed analytically based on full knowledge of the mean responses and the noise. For the SVM-PCA computational observer, which has to learn both the mean responses and the statistics of the noise, performance increases with number of training trials, presumably because the generalizability of the separating hyperplane increases with more data. For the SVM-Template observer, however, whose spatial pooling operation reduces uncertainty regarding the mean responses, performance is relatively stable with the number of trials, consistently about 20% that of the ideal observer inference engine. The SVM-PCA performance exceeds the SVM-Template performance after 8000,1500 and 500 trials for the 8 c/deg 16 c/deg, and 32 c/deg stimuli respectively. So, when the number of trials is high enough for the response dimensionality, SVM-PCA performs better than the SVM-Template engine, since no information is thrown away by the spatial pooling mechanism. Moreover, if we extrapolate the SVM-PCA performance with the number of trials, we approach the ideal observer performance after 16.0, 4.9 and 1.6 million trials, for the 8, 16 and 32 c/deg stimuli, respectively. However, computing very large number of trials of high-dimensionality signals is prohibitive in terms of computational resources. Therefore, given the relative stability of the SVM-Template with respect to the size of the training data set, we decided to employ this classifier for the full simulation in which we compare performance of computational to human observers below.

**Figure 8:**
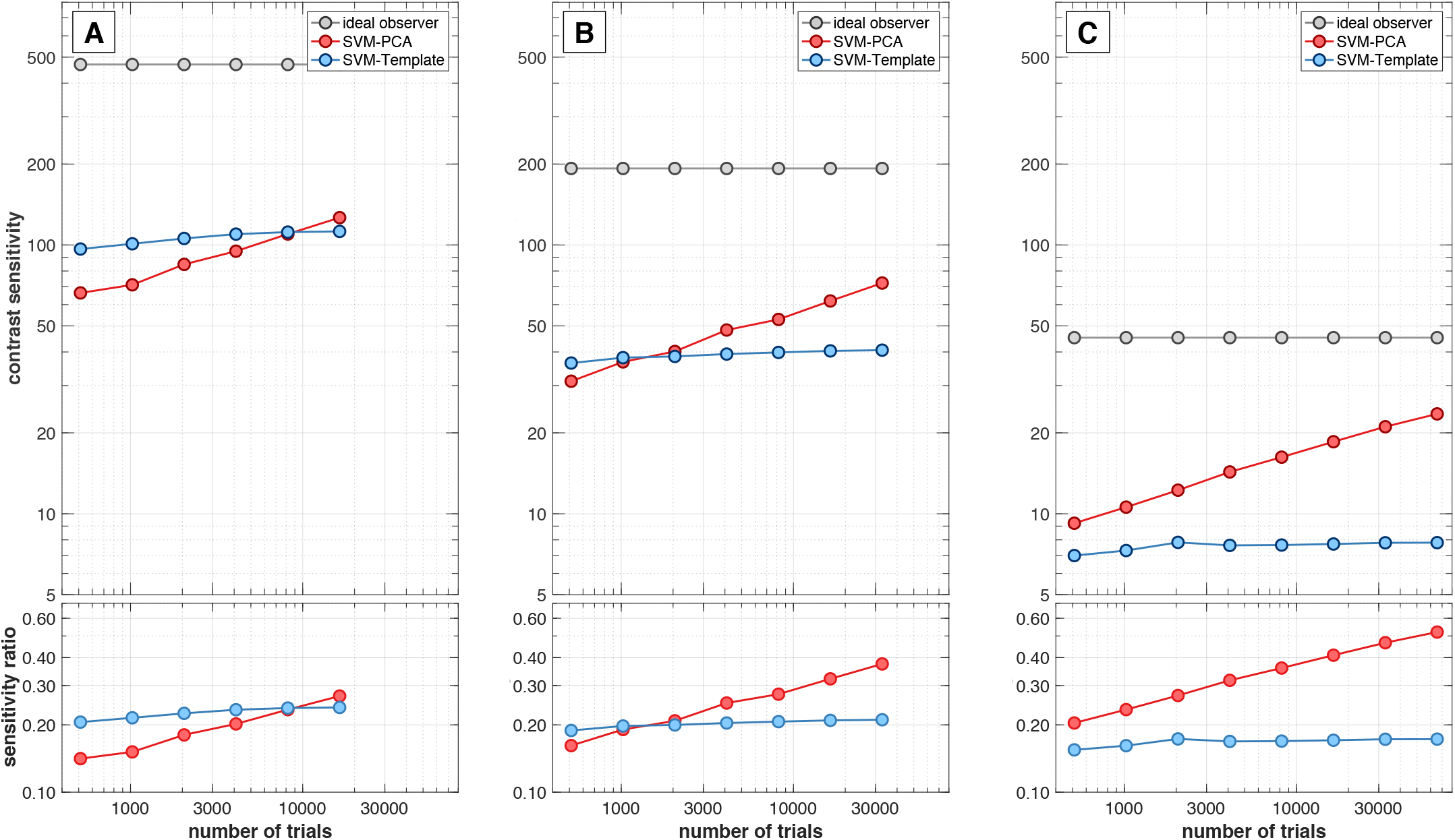
Dependence of computed contrast thresholds on the number of training response instances. **A**. Contrast thresholds at 8 c/deg. **B**. Contrast thresholds at 16 c/deg stimulus. **C**. Contrast thresholds at 32 c/deg stimulus.

### Comparison of computational and human observer performance

Figure 9 compares performance of a full simulation to the data and simulations reported by Banks et al. (1987). All CSFs depicted here were computed for a 2 mm pupil for direct comparison to the human data measured by Banks et al. (1987). The computational-observer CSF shown in blue, was derived used a realistic mosaic (Figure 4D), the optics of our typical Thibos subject (Subject 3, but with PSF computed from the wavefront aberrations for a 2 mm pupil), and the SVM-Template inference engine. This pipeline has a 5-10 fold general sensitivity drop as compared to 1-2 fold sensitivity drop obtained using the ideal observer inference engine (red points), showing the large effect of inference engine on absolute sensitivity. There are modest changes in relative sensitivity between our ideal observer CSF and our replication of the Banks et al. (1987) CSF (gray points). These are a reduction at the lowest spatial frequencies due to the eccentricity-varying cone density, and a slight increase at the highest spatial frequencies (50 and 60 c/deg) due to wavefront-based optics. Overall, the SVM-Template computational observer pipeline brings the computed CSF to within a factor of 2-3 of measurements from the two human observers of Banks et al. (1987).

**Figure 9:**
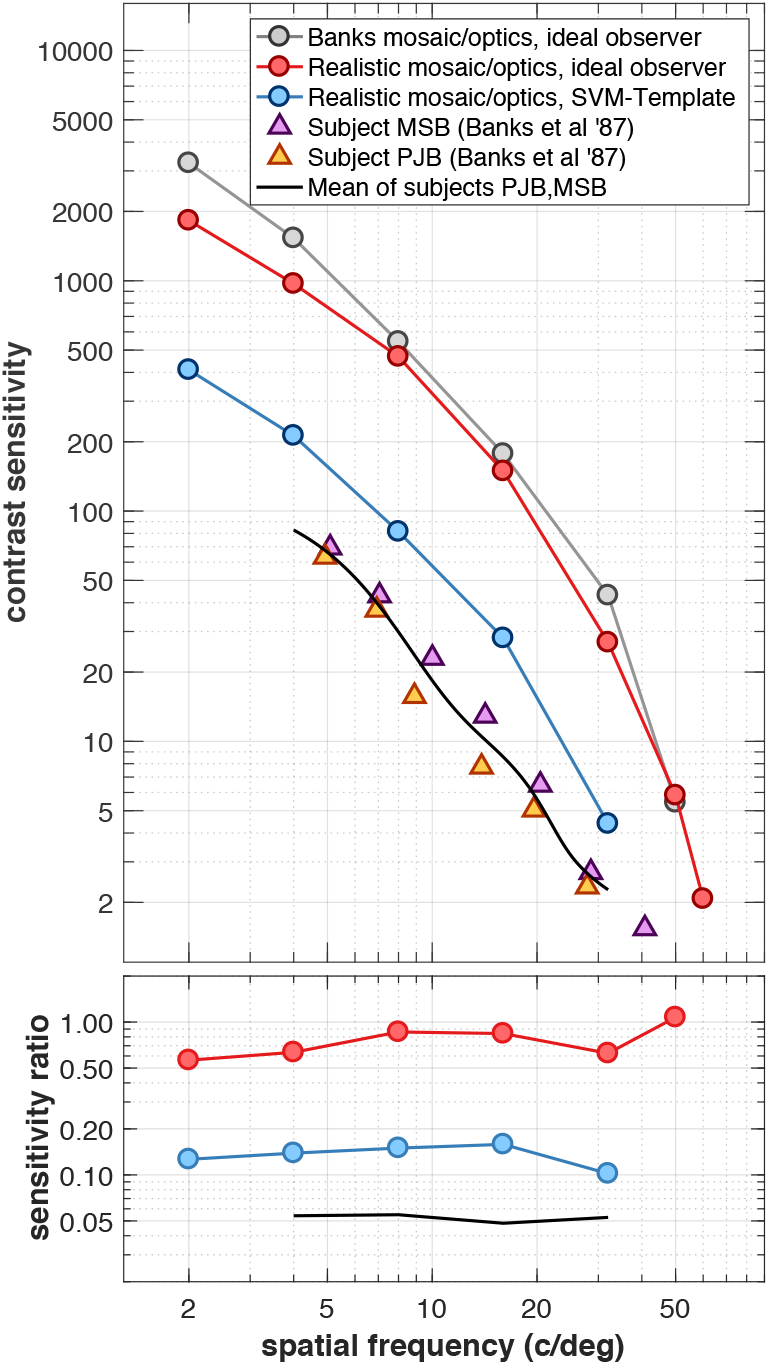
Comparison of computational observer - derived CSFs to CSFs measured in humans. All CSFs are for 2 mm pupils. The ideal observer CSF was derived using the parameters of Banks et al. (1987) and is shown in gray disks (replotted from Figure 3A). The red disks depict the CSF derived using the eccentricity-dependent mosaic (Figure 4D), the typical wavefront-based optics (Figure 5E) and the ideal observer inference engine. It exhibits a modest relative sensitivity decrease at the lowest spatial frequencies, but is otherwise close to that computed by Banks et al. (1987). A 5-fold drop in sensitivity occurs when the inference engine is switched to the data-driven SVM-Template inference engine (blue disks). The CSFs measured in real subjects by Banks et al. (1987) are shown in triangles, and the black line depicts the mean of these subjects’ CSF, estimated by fitting the subject data with a double exponential curve. The CSF measured in human subjects is lower than the SVM-Template CSF by a factor of 2-3.

## Discussion

### The limits of spatial resolution

We applied the ISETBio computational methods to clarify how specific properties of the cone mosaic and the physiological optics limit pattern resolution. First, the spatial structure of the cone mosaic is an important factor in limiting the CSF. The CSF is commonly measured using gratings that cover different amounts of the mosaic, and the change in the cone density across the mosaic is noticable for typical variation in size with frequency. This variation is partly compensated by the change in cone aperture, but even so there remain differences between computations based on uniform and eccentricity-dependent mosaics. Second, modern wavefront measurements indicate better human optics than earlier measurements, and all else equal incorporating the wavefront measurements leads to less attenuation in the ideal observer CSF at high spatial frequencies. Third, the choice of inference engine has a large effect on the absolute level of performance. Certain choices bring the computational observer into much closer agreement with measured performance (SVM linear classifiers). Other choices show that more information is available in principle (e.g., the signal-known-exactly ideal observer). The idea that behavior can be completely described analysis of performance based on the visual representation at the cone mosaic is, of course, wrong. But the ability to calculate the information available to an ideal or computational observer at specific stages of visual processing continues to provide useful benchmarks that clarify the aspects of performance that require explanation in terms of other factors.

### Future directions

We are currently investigating the impact of additional ISETBio computational modules on pattern sensitivity. These include models of fixational eye movements and of the non-linear transformation from cone excitations to photocurrent (Cottaris, Rieke, Wandell, & Brainard, 2018). ISETBio also includes methods based on computer graphics and ray tracing that quantify the retinal images of three-dimensional scenes (Lian, Farrell, & Wandell, 2018). This work can extend ISETBio applications into additional vision science areas, such as accommodation and depth perception.

Retinal and cortical visual processing transforms the cone excitations in many ways that impact visual performance. ISETBio is designed to be extensible, and the current implementation contains placeholders for models of multiple parallel mosaics of retinal bipolar and ganglion cells. For example, understanding the limits of color sensitivity may be accessible through these calculations. Opponent processing of signals from different cone classes is a key step in color coding (Stockman & Brainard, 2010), and quantifying how this combination is implemented in neural circuits remains elusive. Implementing image-computable models of bipolar and ganglion cells is clarifying where there are gaps in our current knowledge of how these cells operate.

### Applications

The enormous growth of the imaging industry is based on the ability to design and implement new optical and electronic devices; during this process designers inevitably turn to vision science for guidance in setting parameters. Critical information includes the standard color observer (Judd & Wyszecki, 1975; Wyszecki & Stiles, 1982), knowledge of pattern resolution (Geisler & Banks, 1995; Wandell, 1995; Watson & Ahumada, 2004, 2005) and position resolution (Westheimer, 1981; Klein & Levi, 1985; Jiang et al., 2017). The computational methods in ISETBio integrate quantitative models of scenes and display devices and are useful in supporting the design and evaluation of new imaging devices.

Medicine is a second important application area. As treatments for partial sight restoration become feasible, for example through gene therapy or retinal prostheses, it will be important to understand the degree to which the additional information provided to the nervous system by these technologies support performance. The ability to use simulations to model the information carried by restored representations and understand the upper limits on visual performance available from them should facilitate the design of therapies that can ameliorate partial and full blindness (Cottaris & Elfar, 2005; Jiang, Wandell, & Farrell, 2015; Golden et al., 2018; Beyeler, Boynton, Fine, & Rokem, 2018).

### Machine learning

Inference-engines with linear SVM-based classifiers bring computational observer performance much closer to human performance levels. Such inference-engines implement a decision variable that is a weighted sum of the representational input (here, cone excitations). Spatial pooling by linear weighted sums is an approximate model for the receptive field properties of several neuronal populations (Movshon, Thompson, & Tolhurst, 1978; Andrews & Pollen, 1979; Shapley, Kaplan, & Soodak, 1981; Wandell, 1995). In addition, the linear classifiers are learned from training data, and do not need to assume that prior knowledge about the stimuli is provided to the visual system. Basing decisions on the weighted sum of cone signals that are learned by linear classifiers may approximate the inference engines used by real observers’ neural processing better than the ideal observer calculation.

Work in machine-learning offers another opportunity for collaboration. Several machine-learning successes use convolutional neural networks to analyze images (Kriegeskorte, 2015). The architecture and parameters of these networks offer inspiration about how to model cortical circuits, and conversely there are opportunities to explore how findings from cortical circuits might be used to implement artificial neural networks (Yamins et al., 2014; Khaligh-Razavi & Kriegeskorte, 2014). A limitation in the interaction between vision science and machine-learning arises from the stimulus representation. Convolutional neural networks are typically trained using digital image values (RGB), which have no biological basis. The machine learning work can be more closely integrated with biology by training on inputs comprising realistic visual inputs. The ISETBio simulations are well-suited for converting RGB images into retinal responses that serve as biologically realistic inputs to train artificial neural networks.

### Theory and computation

Theory is how we develop a principled understanding of complex systems. Computational models built on theoretical principles can provide additional insights about the impact of specific system components and deviations from the ideal. Coordination between theory and computation arises in many fields. Rocket design incorporates Newton’s gravity formulation as well as computational models of material properties, friction and heat. Telecommunications systems incorporate Shannon’s information theory, as well as information about switching times, conduction delays, and circuit noise.

In vision science ideal observer theory informs us how to conceptualize the inputs and decision variables that define system performance. With the enormous growth of computational power, this formal theory - which inevitably involves many approximations - can be extended to account for specific system characteristics. Modeling the impact of these system components is important for bridging basic discovery and applications, say for display engineering or medicine.

## Methods

### Stimulus

The simulated scenes were designed to match the stimuli used by Banks et al. (1987): cosinusoidal patterns windowed using a half-cosine spatial modulation which span 7.5 cycles of the grating. The windowing makes the spatial extent of each stimulus inversely proportional to its spatial frequency, a choice motivated by the observation that (a) contrast sensitivity increases with extent up to a critical size, and (b) the critical size is approximately constant when expressed in terms of stimulus cycles (Howell & Hess, 1978).

ISETBio scenes are spatially sampled spectral radiances (radiometric). The stimuli employed by Banks et al. (1987) are specified in colorimetric units (e.g., luminance). A spectral representation is necessary to model chromatic aberration, inert pigments, and absorption of light by the three classes of cone photoreceptors. To promote the colorimetric specification to spectral radiance, we simulated the scenes as arising from a typical color CRT from the era when their paper was published. The critical display information is the R, G and B channel spectral power distributions, the RGB display quantization, and the pixel spatial sampling. Because our interest here is not the effect of display properties per se, we modeled a CRT with 18-bit linear control of the R, G and B primary intensities. We also set the pixel spatial sampling to be inversely proportional to the stimulus spatial frequency: consequently, all stimuli were represented on a 512x512 spatial grid with this grid mapped onto the corresponding retinal region in a manner that took the stimulus size into account. The spectra we model differ somewhat from those in the Banks et al. (1987) experiment, as the experiment was performed using a monochrome CRT with a P4 phosphor.

### Retinal image

Physiological optics transforms the scene spectral radiance to the retinal image (spectral irradiance). The transformation can be conveniently grouped into two parts. First, the scene radiance is transformed to an idealized retinal spectral irradiance. This transformation accounts for (a) the pupil diameter (which controls the amount of light entering the eye), (b) the stimulus distance and the posterior nodal distance of the lens (which controls the retinal image magnification (Holst, 1989), and (c) the lens pigment spectral transmittance, which reduces retinal irradiance in a wavelength-dependent manner (Stockman, Sharpe, & Fach, 1999). Second, the idealized retinal spectral irradiance is convolved with a wavelength-dependent point spread function (PSF). The PSF is determined by monochromatic and chromatic aberrations of the optics as well as diffraction. Blurring by the wavelength-dependent PSF produces the retinal image.

In general, there are three types of optical aberrations: (i) monochromatic aberrations, (ii) longitudinal chromatic aberration (LCA), and (iii) transverse chromatic aberration (TCA). Monochromatic aberrations produce complex deformations in the retinal image that vary between individuals. LCA is a wavelength-dependent defocus which occurs due to the wavelength-dependent refractive index of the ocular media. LCA is consistent across individuals and amounts to about 2.2 diopters of defocus across the 400-700 nm range of the spectrum (Bedford & Wyszecki, 1957; Thibos et al., 1992; Marimont & Wandell, 1994; Cottaris, 2003). TCA causes a wavelength-dependent shift in the position and magnification of the retinal image. It results from changes in the index of refraction of the optical elements combined with misalignment of these components. TCA varies between individuals and between the eyes of a given individual (Marcos, Burns, Moreno-Barriusop, & Navarro, 1999; Harmening, Tiruveedhula, Roorda, & Sincich, 2012). Because the optical axis of the eye is not always centered with its visual axis, TCA can be observed at the fovea. In some individuals TCA, can be more significant than LCA, whereas in other individuals it can be minimal (Marcos et al., 1999). In the present work we model monochromatic aberrations and LCA. We neglect TCA, as well as changes with wavelength in wave aberrations other than defocus (Marcos et al., 1999). In addition, we neglect light scatter due to the ocular media (Vos, 2003) and the Stiles-Crawford Effect (Stiles & Crawford, 1933; Westheimer, 2008).

We model monochromatic aberrations using the first 15 Zernike polynomials, which were measured in a population of 200 human eyes (Thibos et al., 2002). From a set of Zernike polynomials, we can compute the wavefront aberration map at the in-focus wavelength (550 nm) and from this the corresponding point spread function (Goodman, 2005; Watson, 2015). To generate the point spread function for any other wavelength, we add a defocus term, *d(λ)*, to the Zernike polynomials according to the formula given by Howarth and Bradley (1986):

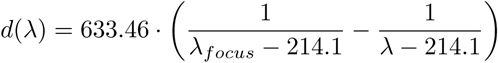

where λ_*focus*_ = 550nm.

#### Selecting representative Thibos subjects

For the CSF simulations we wanted to choose wavefront-based optics for a typical subject (Thibos et al., 1992). As noted previously, using a wavefront function based on the mean of the Zernike coefficients across all subjects is not satisfactory because the cancelation of positive and negative defocus coefficients in the averaging leads to a higher optical MTF than is observed in most subjects. At the same time, deriving typical optics directly from the mean MTF is not straightforward because the MTF does not completely determine the spatial structure of the PSF, so that additional assumptions are required.

To deal with this issue we computed CSFs using optics from 5 sample eyes from the Thibos data set that we choose to span the range of measured optical quality. The PSFs of these subjects are depicted Figure 5, where we use the term subject to refer to a specific eye of a particular subject. The subjects are referred to as Subjects 1, 2, 3, 4 and 5, and we selected Subject 3 as typical. The Zernike coefficients for these 5 subjects are provided in Table 1.

**Table 1:**
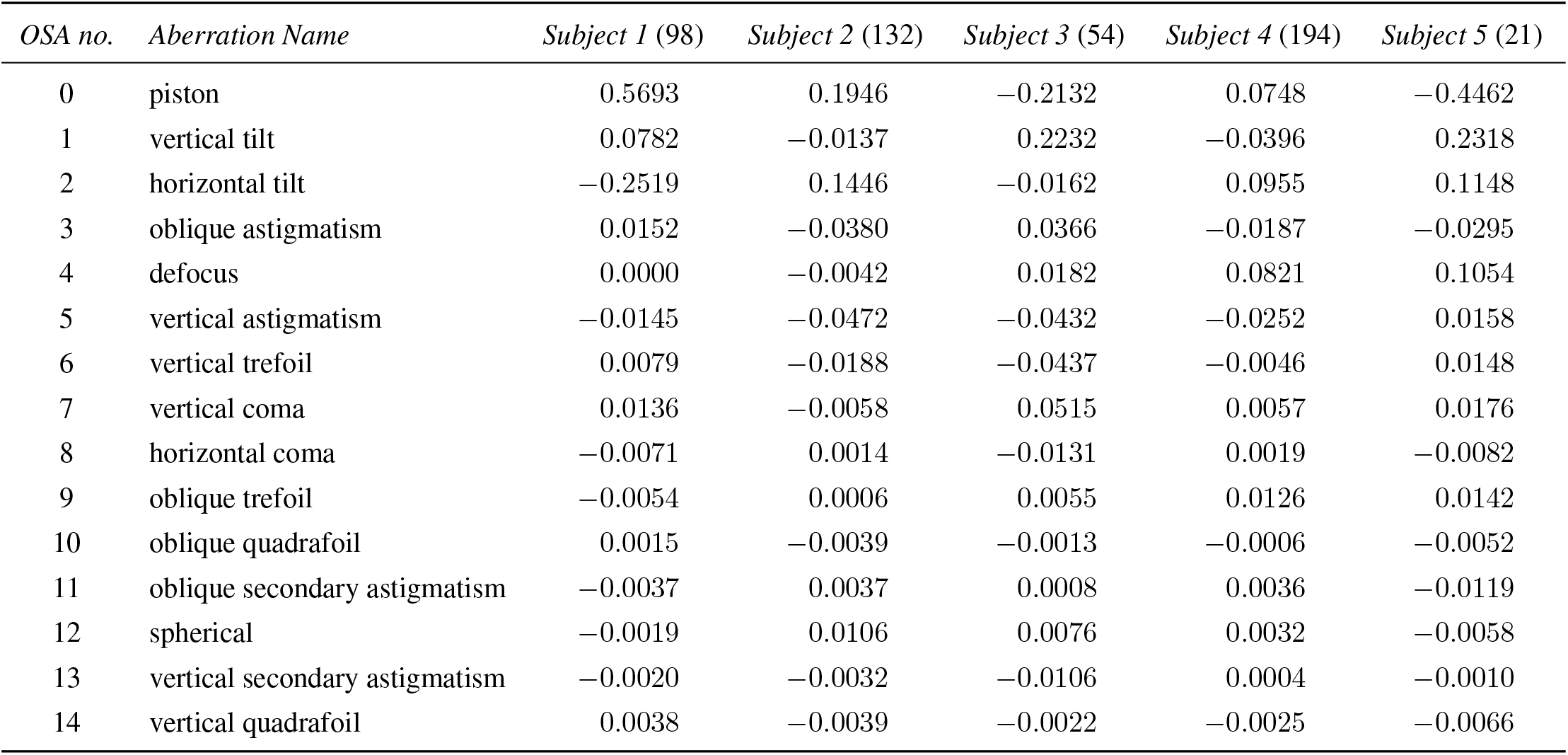
Zernike coefficients for the 5 employed Thibos subjects. These are taken from the 3mm pupil data set of (Thibos et al., 2002). The numbers within the parentheses next to each subject correspond to the index of the ‘OU’ field in the data set, which contains left and right eyes for the population of 100 subjects. Data for 4.5mm, 6 mm and 7.5 mm pupils are available for these subjects in the full (Thibos et al., 2002) dataset.

| OSA no. | Aberration Name | Subject 1 (98) | Subject 2 (132) | Subject 3 (54) | Subject 4 (194) | Subject 5 (21) |
| --- | --- | --- | --- | --- | --- | --- |
| 0 | piston | 0.5693 | 0.1946 | −0.2132 | 0.0748 | −0.4462 |
| 1 | vertical tilt | 0.0782 | −0.0137 | 0.2232 | −0.0396 | 0.2318 |
| 2 | horizontal tilt | −0.2519 | 0.1446 | −0.0162 | 0.0955 | 0.1148 |
| 3 | oblique astigmatism | 0.0152 | −0.0380 | 0.0366 | −0.0187 | −0.0295 |
| 4 | defocus | 0.0000 | −0.0042 | 0.0182 | 0.0821 | 0.1054 |
| 5 | vertical astigmatism | −0.0145 | −0.0472 | −0.0432 | −0.0252 | 0.0158 |
| 6 | vertical trefoil | 0.0079 | −0.0188 | −0.0437 | −0.0046 | 0.0148 |
| 7 | vertical coma | 0.0136 | −0.0058 | 0.0515 | 0.0057 | 0.0176 |
| 8 | horizontal coma | −0.0071 | 0.0014 | −0.0131 | 0.0019 | −0.0082 |
| 9 | oblique trefoil | −0.0054 | 0.0006 | 0.0055 | 0.0126 | 0.0142 |
| 10 | oblique quadrafoil | 0.0015 | −0.0039 | −0.0013 | −0.0006 | −0.0052 |
| 11 | oblique secondary astigmatism | −0.0037 | 0.0037 | 0.0008 | 0.0036 | −0.0119 |
| 12 | spherical | −0.0019 | 0.0106 | 0.0076 | 0.0032 | −0.0058 |
| 13 | vertical secondary astigmatism | −0.0020 | −0.0032 | −0.0106 | 0.0004 | −0.0010 |
| 14 | vertical quadrafoil | 0.0038 | −0.0039 | −0.0022 | −0.0025 | −0.0066 |

We selected the subjects by ranking the entire population of 200 eyes measured by Thibos et al. (1992) and by chosing 5 subjects whose scores span the range of computed scores. Subject ranking was done as follows. First, we computed the singular value decomposition of all subject PSFs, separately for each wavelength. This provided a basis set for representing the PSFs at each wavelength. We then projected each subject’s PSF and the mean Zernike coefficient PSF to the basis set, separately for each wavelength. A PSF matching score was computed for each subject based on the mean (over wavelengths) RMS error between that subject’s projection coefficients and the projection coefficients of the mean Zernike coefficient PSF. Then, we computed the mean MTF (absolute value of the OTF) across all subjects. An MTF matching score was computed for each subject as the mean (over wavelengths) RMS error between that subject’s MTF and the mean MTF. The calculations were done for a 3 mm pupil. Subjects were ranked according to their PSF score (Figure 10), and the 5 representative subjects were selected as follows. Subject 1 was selected because his or her PSF best resembled the PSF obtained from the mean of the Zernike coefficients. Note that this subject’s MTF score is very low. Subject 2 also has a high PSF score but a much higher MTF score. Subject 3 has PSF and MTF scores of similar magnitude. This is the subject we take to represent typical human optics. Subjects 4 and 5 have progressively worse PSF scores and low MTF (Figure 10).

**Figure 10:**
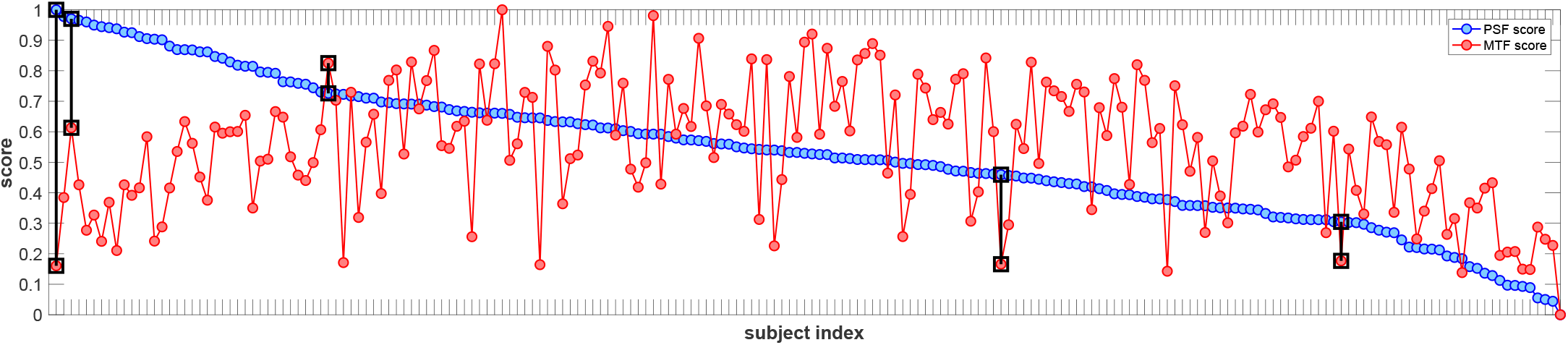
Selecting representative subjects. PSF and MTF matching scores for the population of the 200 Thibos subjects sorted according to their PSF score. The 5 representative subjects are indicated by the black squares which connect the subjects’ PSF and MTF scores, with subjects 1-5 running from left to right in the figure.

### Computation of cone excitations (photon isomerizations)

The main factors that determine how the retinal image, *RI*(*x,y,λ*), is transformed into a pattern of cone photoisomerization rates are (i) the macular pigment, which differentially absorbs short wavelength photons, (ii) the spectral quantal efficiency, or spectral absorptance, of the cone photopigment, which controls the proportion of incident photons that get absorbed by the photopigment, (iii) the cone aperture diameter, which determines the photon collecting area of a cone and which also acts as a spatial low-pass filter, and (iv) the cone lattice, which controls the spatial sampling of the retinal irradiance image.

The macular pigment transmittance, *T_macular_*(λ), is depicted in Figure 11A. Minimum transmittance is 0.45 at 460 nm. In our calculations, we do not model the variation of *T_macular_*(λ) with eccentricity (Bone, Landrum, Fernandez, & Tarsis, 1988). The spectral quantal efficiencies (absorptances) of different cone classes, *q_c_*(λ), with *c* = {L,M,S}, are depicted in Figure 11B and are computed based on the Stockman-Sharpe normalized absorbance values, *SS_c_*(λ), (Stockman et al., 1999; Stockman & Sharpe, 2000), as:

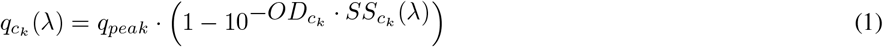

where *q_peak_* is the peak cone quantal efficiency, 0.667 for all cone types, and *OD_ck_* is optical density of cone type *c_k_*, 0.5 for L- and M-cones and 0.4 for S cones. These values are within the range of optical densities reported, 0.29 – 0.91 for L-cones, 0.36 – 0.97 for M-cones (Renner, Knau, Neitz, Neitz, & Werner, 2004).

Cones exhibit waveguide properties (Enoch, 1961), according to which light incident on the cone inner segment is guided to the outer segment, where it gets absorbed. To model this, we employed a spatially uniformly-weighted circular averaging filter, *A*(*x,y*), whose diameter corresponds to the inner segment diameter and whose volume is 1. In the Banks et al. (1987) mosaic, the inner segment diameter is 3 *μ*m, whereas in the ISETbio mosaics, it is 1.6 *μ*m in the fovea. The aperture filters and the corresponding modulation transfer functions for these mosaics are depicted in Figure 11C. Note that the modulation transfer function at 60 cycles/deg is 0.63 for the mosaic employed by Banks et al. (1987) mosaic and 0.89 for the eccentricity-varying cone mosaics. Although we varied the size of the inner segment diameter with eccentricity in when we computed cone excitations, we used a constant inner segment diameter, taken as the value at the fovea, when computing blur by the cone apertures. This choice was made for reasons of computational efficiency.

**Figure 11:**
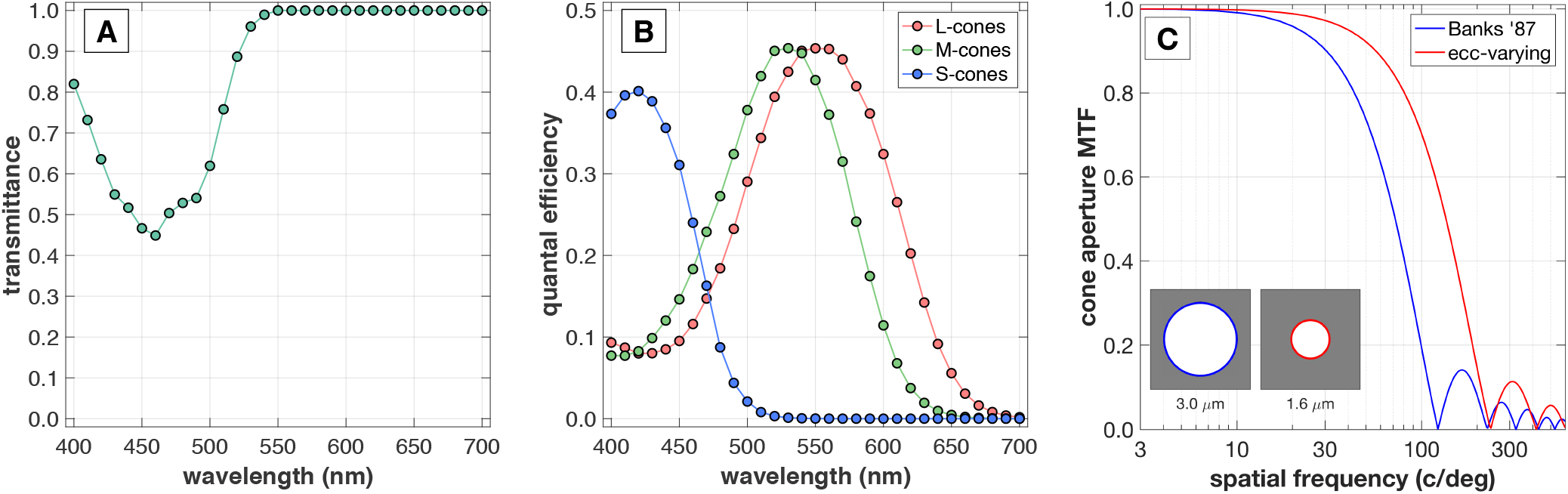
Components of the cone excitation response model. **A**. Macular pigment transmittance as a function of wavelength, *Tmacuiar*(λ). **B**. Spectral quantal efficiencies, *q_c_*(λ), for *c* = *L,M,S* cones. **C**. Cone aperture filters (inset) and corresponding MTFs for the foveal inner segment diameters considered in this paper.

To compute the spatial distribution of cone excitation rate, *CER_c_(x,y)*, for each cone class, *c*, the retinal image, *RI(x, y, λ)*, was first filtered with the the macular pigment transmittance, *T_macular_*(λ). The for each cone class we multiplied by the corresponding spectral quantal efficiency, *q_c_*(λ) and integrated over wavelength. The result was then spatially convolved with the cone aperture, *A*(*x, y*):

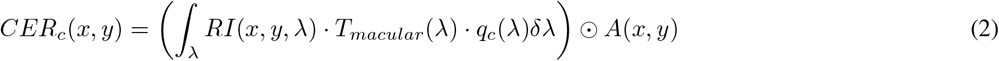

To compute the cone mosaic excitation we estimate the mean count of cone excitation events within the simulation time interval, here τ = 5 msec. Specifically, for a cone *k*, of class *c_k_* located at coordinates (*x_k_,y_k_*), the mean count of cone excitation events, 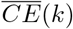, within τ msec is computed by spatially-sampling the continuous function *CER_ck_*(*x,y*) at (*x = x_k_, y = y_k_*), multiplying by the cone inner segment area, *α*, and by the time interval, *τ*:

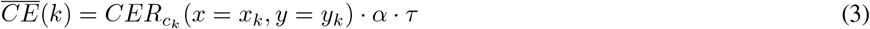

Finally, an excitation response instance, *i*, for the *k*-th cone, *CE^i^*(*k*), is generated by adding a random number of excitation events drawn from a Poisson distribution whose mean is equal to the mean count of excitation events:

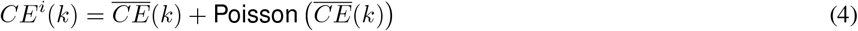

#### Eccentricity-dependent cone efficiency correction

The above computation or isomerization does not take into account the fact that as eccentricity increases, inner segment area increases (Curcio et al., 1990), whereas outer-segment length decreases (Banks, Sekuler, & Anderson, 1991; Jonnal et al., 2017). We approximated these effects by defining an eccentricity dependent correction factor, *b_k_*, for the k-th cone located at (*x_k_, y_k_*) defined as:

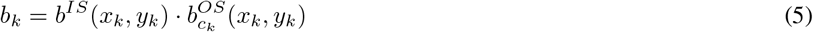

where

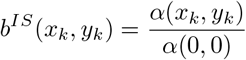

is the correction factor required to account for the change in inner segment area, *α*(*x_k_, y_k_*), at location (*x_k_, y_k_*), relative to its foveal value *α*(0,0). The quantity 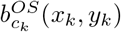 is the correction factor required to account for the decrease in outer segment length for cone class *c_k_* at location (*x_k_, y_k_*), relative to its foveal value, and is computed as the mean value of 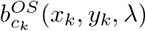 over the wavelength parameter λ, with

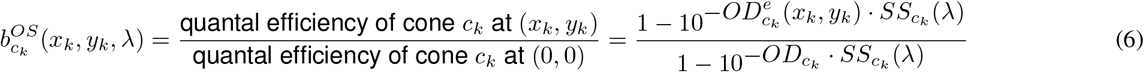

with

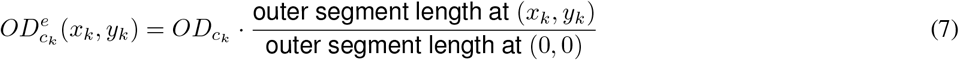

Ideally, 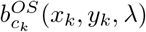 should be applied within the integral of Equation 2. For ease of computation, we use the mean value over all wavelengths of 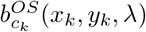 and apply the *b_k_* correction factor (Equation 5) to the mean count of excitation events, 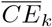, to update the quantity computed by equation 3:

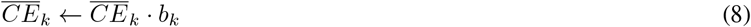

This allows us to compute a cone excitation count which takes into account the eccentricity-dependent changes in cone efficiency due to changes in inner segment aperture and outer segment length. In these computations, we assume that photopigment concentration and extinction coefficients remain constant across eccentricity, and we ignore photopigment bleaching which is small at these light levels (Rushton & Henry, 1968).

### Cone mosaics

We examined two types of hexagonal cone mosaics, (i) the regularly-spaced hexagonal mosaic employed by Banks et al. (1987), in which cone density is constant across all eccentricities with a spacing of 3 *μ*m and an inner segment diameter of 3 *μ*m, and (ii) eccentricity-dependent mosaics, in which cone density varies with eccentricity as described in Curcio et al. (1990). In eccentricity-dependent mosaics, the minimum cone spacing at the foveola is 2 *μ*m. This corresponds to a peak cone density of 250,000 cones/mm^2^ (Curcio et al., 1990). The ratio of cone diameter to inner segment aperture ratio is 0.79, close to the 0.82 value suggested by (Miller & Bernard, 1983; Curcio et al., 1990), across all eccentricities.

The eccentricity-dependent lattices are constructed using a three-step process. First, a regular hexagonal lattice is constructed with the minimum (foveal) cone separation (Figure 12A). Next, the regular hexagonal lattice is subsampled using an eccentricity-dependent probability of eliminating cones. The function is chosen so that the expected density at each eccentricity matches a target density. The resulting spatial mosaic approximates the desired eccentricity-dependent cone density but the resulting cone coverage is non-uniform with clumps of cones in some locations separated by regions without any cones (Figure 12B). Subsequently, cone positions are adjusted to improve the local uniformity using an iterative procedure (Persson, 2005; Figures 12C-E). In this approach, a cone and its neighboring cones are subjected to simulated movement driven by mutually repulsive forces. Finally, cones are assigned a type, L, M, or S, depending on the specified L/M/S cone density property, as well as the desired S-cone mosaic properties, such as a minimum distance between neighboring S-cones and the size of an S-cone free central region (Figure 12F).

**Figure 12:**
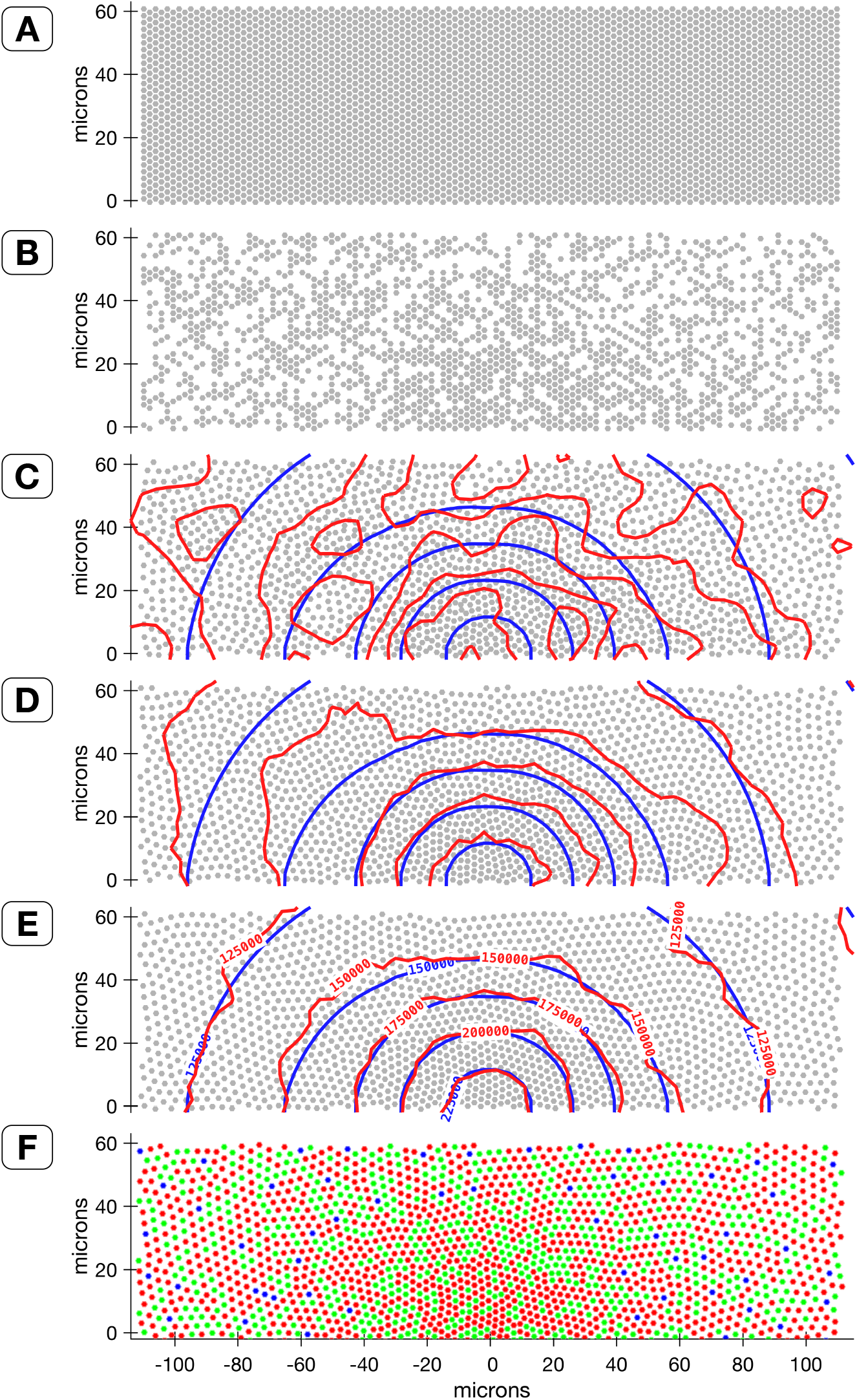
Procedure for generation of approximately-hexagonal mosaics with eccentricity-varying density. **A**. The mosaic is initialized with a regular hexagonal lattice of the minimum cone spacing. **B**. Next, the mosaic is probabilistically subsampled based on cone density data from Curcio et al. (1990). **C.-E**. Subsequently, cone positions are iteratively adjusted while trying to maintain proximity of the achieved cone density map (red contours) to the desired cone density map (blue contours). **F**. Finally, cones are assigned types, L, M, or S, depicted as red, green and blue symbols, according to the specified mosaic properties.

### Inference engine

An ideal inference engine for cone excitations modeled as Poisson processes is constructed from knowledge of the mean isomerization counts to the test and null stimuli. This signal-known-exactly calculation defines an upper bound on the information that can be extracted. For more realistic calculations, say accounting for uncontrolled fixational eye movements, the cone excitations have additional uncertainty and across trials the noise is no longer Poisson. With these additional terms, a simple closed form mathematical expression describing the cone excitation signals across trials may be beyond our reach.

For the general case it is possible to choose an inference-engine that learns a linear classifier from training examples. In this study we employ Support Vector Machines (SVM) (Scholkopf & Smola, 2002; Manning et al., 2008) that learn a linear classifier (Figure 6). A general challenge in the implementation concerns the high dimensionality of the cone excitations. In this study, for example, the smallest cone mosaic had 5,640 dimensions (20 time bins × 282 cones) and the largest cone mosaic had 1,059,380 dimensions (20 time bins × 52,969 cones). To train inference-engines based on SVM linear classifiers, we start with dimensionality reduction. In the present study, we compared two different techniques.

*SVM-PCA classifier.* For the first dimensionality reduction technique, we computed the first 60 principal components of a data set comprising 1024 examples of the null and test cone responses. The PCA was performed separately for each stimulus condition. Binary SVM classification is performed on the projections of the response instances into the space spanned by the 60 PCAs. We refer to this classifier as the SVM-PCA classifier.

*SVM-Template classifier.* The second dimensionality reduction technique uses spatial pooling via a weighting kernel, or template, *V*(*k*), *k* = 1… *M*, where *M* is the number of cones in the mosaic. The spatial profile of *V(k)* is derived from the spatial contrast modulation of the test stimulus (Figure 13). Given a set of null and test stimulus response instance vectors, 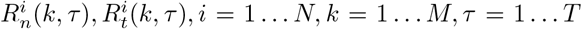, we begin by computing the mean, over the *N* instances and *T* time bins, response of each cone, *k*, to the null stimulus, 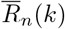. This mean response to the null stimulus is subtracted from both the test and the null response instance vectors, and the inner product between the mean-subtracted cone excitations and the template is computed to simulate spatial pooling using the *V*(*k*) template:

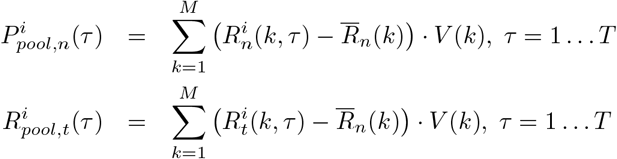

**Figure 13:**
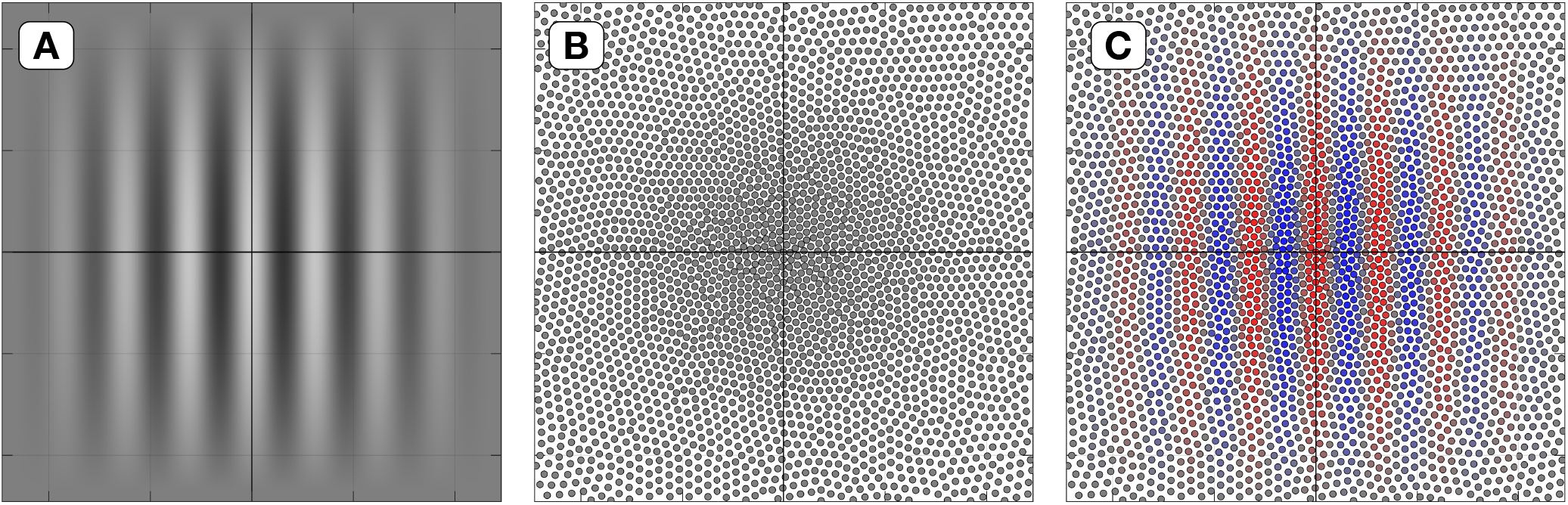
Stimulus-matched spatial pooling template. **A**. The spatial contrast modulation for the 16 cycles/deg stimulus **B**. The employed cone mosaic. **C**. The spatial pooling kernel, or template, for this stimulus and this mosaic with which cone responses are weighted before pooled. Red indicates positive weights, blue indicates negative weights, whereas saturation indicates weight strength.

The spatially-pooled response instance vectors 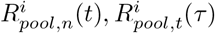 are used to train the SVM linear classifier. We refer to this classifier as the SVM-Template classifier. In the present paper, in which we concentrate on cone excitation responses, *R^i^*(*k,τ*) is the quantity *CE^i^*(*k,τ*) (Equation 4).

## References

Andrews, B. W., & Pollen, D. A. (1979). Relationship between spatial-frequency selectivity and receptive-field profile of simple cells. The Journal of Physiology, 287, 163–176. [PubMed]

Angueyra, J., & Rieke, F. (2013). Origin and impact of phototransduction noise in primate cone photoreceptors. Nature Neuroscience, 16(11), 1692–700. [PubMed] [Article]

Artal, P. (2015). Image formation in the living human eye. Annual Review of Vision Science, 1(1), 1–17. [PubMed] [Article]

Banks, M., Geisler, W., & Bennett, P. (1987). The physical limits of grating visibility. Vision Research, 27 (11), 1915–1924. [PubMed] [Article]

Banks, M., Sekuler, A., & Anderson, S. (1991). Peripheral spatial vision: limits imposed by optics, photoreceptors, and receptor pooling. Journal of the Optical Society of America. A, 8, 1775–1787. [PubMed]

Barlow, H. (1964). The Physical Limits Of Visual Discrimination (Vol. 2; A. C. Giese, Ed.). New York: Academic Press.

Baylor, D., Nunn, B., & Schnapf, J. (1984). The photocurrent, noise and spectral sensitivity of rods of the monkey macaca fascicularis. The Journal of Physiology, 357, 575–607. [PubMed] [Article]

Bedford, R. E., & Wyszecki, G. (1957). Axial chromatic aberration of the human eye. Journal of the Optical Society of America. A, 47 (6), 564–565. [Article]

Beyeler, M., Boynton, G. M., Fine, I., & Rokem, A. (2018). pulse2percept: A python-based simulation framework for bionic vision. bioRxiv 148015; doi: https://doi.org/10.1101/148015.

Bone, R. A., Landrum, J. T., Fernandez, L., & Tarsis, S. L. (1988). Analysis of the macular pigment by hplc: retinal distribution and age study. Investigative Ophthalmology & Visual Science, 29(6), 843. [PubMed] [Article]

Bowmaker, J., Dartnall, H., & Mollon, J. (1980). Microspectrophotometric demonstration of four classes of photoreceptor in an old world primate, macaca fascicularis. Nature Neuroscience, 298, 131–43. [PubMed] [Article]

Brindley, G. (1960). Physiology Of The Retina And The Visual Pathway. London: Arnold.

Campbell, F. W., & Gubisch, R. W. (1966). Optical quality of the human eye. The Journal of Physiology, 186(3), 558–78. [PubMed]

Campbell, F. W., & Robson, J. G. (1968). Application of fourier analysis to the visibility of gratings. The Journal of Physiology, 197 (3), 551–566. [PubMed] [Article]

Cottaris, N. P. (2003). Artifacts in spatiochromatic stimuli due to variations in preretinal absorption and axial chromatic aberration: implications for color physiology. Journal of the Optical Society of America. A, 20(9), 1694–1713. [Article]

Cottaris, N. P., & Elfar, S. D. (2005). How the retinal network reacts to epiretinal stimulation to form the prosthetic visual input to the cortex. Journal of Neural Engineering, 2(1), S64–90. [PubMed] [Article]

Cottaris, N. P., Rieke, F. W., Wandell, B. A., & Brainard, D. (2018). Computational observer modeling of the limits of human pattern resolution. In OSA Fall Vision Meeting Abstract.

Curcio, C. A., Sloan, K. R., Kalina, R. E., & Hendrickson, A. E. (1990). Human photoreceptor topography. The Journal of Comparative Neurology, 292(4), 497–523. [PubMed] [Article]

De Vries, H. (1943). The quantum character of light and its bearing upon the threshold of vision, the differential sensitivity and acuity of the eye. Physica, 10(7), 553–564. [Article]

Engbert, R., & Kliegl, R. (2004). Microsaccades keep the eyes’ balance during fixation. Psychological science, 15, 431–6. [PubMed]

Enoch, J. M. (1961). Nature of the transmission of energy in the retinal receptors. Journal of the Optical Society of America. A, 51(10), 1122–1126. [PubMed] [Article]

Farrell, J. E., Jiang, H., Winawer, J., Brainard, D. H., & Wandell, B. A. (2014). Modeling visible differences: The computational observer model. SID Symposium Digest of Technical Papers, 45(1), 352–356. [Article]

Geisler, W. S. (1984). Physical limits of acuity and hyperacuity. Journal of the Optical Society of America. A, 1(7), 775–782. [PubMed]

Geisler, W. S. (1989). Sequential ideal-observer analysis of visual discriminations. Psychological Review, 96(2), 267–314. [PubMed]

Geisler, W. S., & Banks, M. S. (1995). Visual performance. In M. Bass (Ed.), Handbook of Optics: Volume 1. Fundamentals, Techniques, And Design. (p. 1–55). New York: McGraw Hill.

Golden, J. R., Erickson-Davis, C., Cottaris, N., Parthasarathy, N., Rieke, F., Brainard, D. H., et al. (2018). Simulation of visual perception and learning with a retinal prosthesis. bioRxiv 206409, https://doi.org/10.1101/206409.

Goodman, J. W. (2005). Introduction To Fourier Optics (3rd ed.). Roberts & Co: Academic Press.

Harmening, W. M., Tiruveedhula, P., Roorda, A., & Sincich, L. C. (2012). Measurement and correction of transverse chromatic offsets for multi-wavelength retinal microscopy in the living eye. Biomedical Optics Express, 3(9), 2066–2077. [Article]

Holst, G. C. (1989). CCD Arrays, Cameras And Displays, 2nd edition.

Howarth, P. A., & Bradley, A. (1986). The longitudinal chromatic aberration of the human eye, and its correction. Vision Research, 26(2), 361–366. [PubMed] [Article]

Howell, E., & Hess, R. (1978). The functional area for summation to threshold for sinusoidal gratings. Vision Research, 18(4), 369–374. [PubMed] [Article]

Jiang, H., Cottaris, N. P., Golden, J., Brainard, D. H., Farrell, J. E., & Wandell, B. A. (2017). Simulating retinal encoding: factors influencing vernier acuity. Electronic Imaging, Human Vision and Electronic Imaging, 177–181. [Article]

Jiang, H., Wandell, B. A., & Farrell, J. E. (2015). D-CIELAB: A color metric for dichromatic observers. In SID Symposium Digest of Technical Papers. Vol. 46., No. 1.

Jonnal, R. S., Iwona, G., Migacz, J. V., Mehdi, A., Zawadzki, R. J., & Werner, J. S. (2017). The properties of outer retinal band three investigated with adaptive-optics optical coherence tomography. Investigative Ophthalmology and Visual Science, 58(11), 4559–4568. [Article]

Judd, D., & Wyszecki, G. (1975). Color In Business, Science, And Industry. New York: John Wiley and Sons.

Khaligh-Razavi, S., & Kriegeskorte, N. (2014). Deep supervised, but not unsupervised, models may explain it cortical representation. PLoS Computational Biology, 10(11). [Article]

Kingdom, F., & Prins, N. (2010). Psychophysics: A Practical Introduction. San Diego, CA: Academic Press.

Klein, S. A., & Levi, D. M. (1985). Hyperacuity thresholds of 1 sec: theoretical predictions and empirical validation. Journal of the Optical Society of America. A, 2(7), 1170–1190. [Article]

Kriegeskorte, N. (2015). Deep neural networks: A new framework for modeling biological vision and brain information processing. Annual Review of Vision Science, 1(1), 417–446. [Article]

Li, P., Field, G., Greschner, M., Ahn, D., Gunning, D., Mathieson, K., et al. (2014). Retinal representation of the elementary visual signal. Neuron, 81(1), 130–139. [PubMed] [Article]

Lian, T., Farrell, J., & Wandell, B. A. (2018). Image systems simulation for 360 camera rigs. In IST Electronic Imaging Conference, San Francisco.

Liang, J., & Williams, D. (1997). Aberration and retinal image quality of the normal human eye.Journal of the Optical Society of America. A, 14, 2873–83. [PubMed]

Lopez, H., Murray, H., & Goodenough, D. (1992). Objective analysis of ultrasound images by use of a computational observer. IEEE Transactions on Medical Imaging, 11, 496–506.

Manning, C. D., Raghavean, P., & Schutze, H. (2008). Introduction To Information Retrieval. Cambridge: Cambridge University Press.

Marcos, S., Burns, S. A., Moreno-Barriusop, E., & Navarro, R. (1999). A new approach to the study of ocular chromatic aberrations. Vision Research, 39(26), 4309–23. [PubMed]

Marimont, D. H., & Wandell, B. A. (1994). Matching color images: the effects of axial chromatic aberration. Journal of the Optical Society of America. A, 11(12), 3113–3122. [Article]

Martinez-Conde, S., Macknik, S. L., & Hubel, D. H. (2004). The role of fixational eye movements in visual perception. Nature Reviews Neuroscience, 5(229). [PubMed]

Meister, M., & Berry, M. J. (1999). The neural code of the retina. Neuron, 22(3), 435–450. [PubMed]

Miller, W. H., & Bernard, G. D. (1983). Averaging over the foveal receptor aperture curtails aliasing. Vision Research, 23(12), 1365–1369. [PubMed] [Article]

Movshon, J., Thompson, I., & Tolhurst, D. (1978). Spatial summation in the receptive fields of simple cell’s in the cat’s striate cortex. The Journal of Physiology, 53–77.

Pelli, D. G. (1990). The quantum efficiency of vision. In C. Blakemore (Ed.), Vision: Coding And Efficiency (p. 324). Cambridge: Cambridge University Press.

Persson, P. (2005). Mesh generation for implicit geometries. PhD dissertation, MIT.

Pharr, M., & Humphreys, G. (2010). Physically Based Rendering: From Theory To Implementation (2nd ed.). San Francisco: Morgan Kaufmann Publishers.

Pugh, J., E.N., & Lamb, T. (2000). Phototransduction in vertebrate rods and cones: molecular mechanisms of amplification, recovery and light adaptation. In D. Stavenga, W. de Grip, & E. Pugh (eds.), Handbook of Biological Physics, Vol. 3, Molecular Mechanisms of Visual Transduction (p. 183–255). Amsterdam: Elsevier.

Renner, A. B., Knau, H., Neitz, M., Neitz, J., & Werner, J. S. (2004). Photopigment optical density of the human foveola and a paradoxical senescent increase outside the fovea. Visual neuroscience, 21(6), 827–34. [PubMed]

Robson, J. G. (1966). Spatial and temporal contrast sensitivity functions of the visual system. Journal of the Optical Society of America. A, 56, 1141–1142.

Rodieck, R. (1998). The First Steps In Seeing. Sunderland,Mass:Sinauer.

Rose, A. (1948). The sensitivity performance of the human eye on an absolute scale. Journal of the Optical Society of America. A, 38(2), 196–208. [PubMed] [Article]

Rushton, W., & Henry, G. (1968). Bleaching and regeneration of cone pigments in man. Vision Research, 8(6), 617–631. [Article]

Scholkopf, B., & Smola, A. (2002). Learning With Kernels. Cambridge, MA: MIT Press.

Shapley, R., Kaplan, E., & Soodak, R. (1981). Spatial summation and contrast sensitivity of x and y cells in the lateral geniculate nucleus of the macaque. Nature, 292. [Article]

Stiles, W., & Crawford, B. (1933). The luminous efficiency of rays entering the eye pupil at different points. Proceedings of the Royal Society of London B: Biological Sciences, 112(778), 428–450. [Article]

Stockman, A., & Brainard, D. (2010). Color vision mechanisms. In M. Bass, C. DeCusatis, & J. Enoch (Eds.), The Optical Society of America Handbook of Optics, Volume: 3, Vision and Vision Optics (p. 1.11–11.104). New York: McGraw Hill.

Stockman, A., & Sharpe, L. T. (2000). Spectral sensitivities of the middle-and long-wavelength sensitive cones derived from measurements in observers of known genotype. Vision Research, 40, 1711–1737. [PubMed]

Stockman, A., Sharpe, L. T., & Fach, C. C. (1999). The spectral sensitivity of the human short-wavelength cones. Vision Research, 39, 2901–2927. [PubMed]

Tanner, W. J., & Swets, J. (1954). The human use of information I. signal detection for the case of a signal known exactly. Transactions of the IRE Profession Group in Information Theory, 4(4), 213–221. [Article]

Thibos, L. N., Hong, X., Bradley, A., & Cheng, X. (2002). Statistical variation of aberration structure and image quality in a normal population of healthy eyes. Journal of the Optical Society of America. A, 19(12), 2329–2348. [PubMed] [Article]

Thibos, L. N., Ye, M., Zhang, X., & Bradley, A. (1992). The chromatic eye: a new reduced-eye model of ocular chromatic aberration in humans. Applied Optics, 31(19), 3594–3600. [Article]

Tuten, W. S., Cooper, R. F., Tiruveedhula, P., Dubra, A., Roorda, A., Cottaris, N. P., et al. (2018). Spatial summation in the human fovea: the effect of optical aberrations and fixational eye movements. In Press, Journal of Vision. Preprint available at https://doi.org/10.1101/283119.

Vos, J. J. (2003). On the cause of disability glare and its dependence on glare angle, age and ocular pigmentation. Clinical and Experimental Optometry, 86(6), 363–370. [Article]

Wandell, B. A. (1995). Foundations Of Vision. Sunderland, MA: Sinauer.

Watson, A. (2015). Computing human optical point spread functions. Journal of Vision, 15(2), 26.

Watson, A., & Ahumada, A. (2004). The spatial standard observer. Journal of Vision, 4(8), 51–51.

Watson, A., & Ahumada, A. (2005). A standard model for foveal detection of spatial contrast. Journal of Vision, 5(9), 6. [Article]

Watson, A., & Yellott. (2012). A unified formula for light-adapted pupil size. Journal of Vision,12(10), 12. [PubMed]

Westheimer, G. (1981). Visual hyperacuity. In Progress In Sensory Physiology (p. 1–30). Berlin, Heidelberg: Springer.

Westheimer, G. (2008). Directional sensitivity of the retina: 75 years of Stiles–Crawford effect. Proceedings of the Royal Society of London B: Biological Sciences, 275(1653), 2777–2786. [Article]

Westheimer, G., & Campbell, F. W. (1962). Light distribution in the image formed by the living human eye. Journal of the Optical Society of America. A, 52(9), 1040–1045. [PubMed] [Article]

Williams, D. (1985). Visibility of interference fringes near the resolution limit. Journal of the Optical Society of America. A, 2(7), 1087–1093. [PubMed] [Article]

Wyszecki, G., & Stiles, W. S. (1982). Color Science: Concepts And Methods, Quantitative Data And Formulas.

Yamins, D. L. K., Hong, H., Cadieu, C. F., Solomon, E. A., Seibert, D., & DiCarlo, J. J. (2014). Performance-optimized hierarchical models predict neural responses in higher visual cortex. Proceedings of the National Academy of Sciences, 111(23), 8619–8624. [Article]

